# Mapping QTL for vernalization requirement identified adaptive divergence of the candidate gene *Flowering Locus C* in polyploid *Camelina sativa*

**DOI:** 10.1101/2023.05.23.541983

**Authors:** Raju Chaudhary, Erin E. Higgins, Christina Eynck, Andrew G. Sharpe, Isobel A. P. Parkin

**Author notes:** Corresponding author: Isobel Parkin.

## Abstract

Vernalization requirement is an integral component of flowering in winter-type plants. The availability of winter ecotypes among *Camelina* species facilitated the mapping of QTL for vernalization requirement in *C. sativa*. An inter- and intraspecific crossing scheme between related *Camelina* species, where two different sources of the winter-type habit were used, resulted in the development of two segregating populations. Linkage maps generated with sequence-based markers identified three QTL associated with vernalization requirement in *C. sativa*; two from the inter-specific (chromosomes 13 and 20) and one from the intra-specific cross (chromosome 8). Notably, the three loci were mapped to different homologous regions of the hexaploid *C. sativa* genome. All three QTL were found in proximity to *FLOWERING LOCUS C* (*FLC*), variants of which have been reported to affect the vernalization requirement in plants. Temporal transcriptome analysis for winter-type *Camelina alyssum* demonstrated reduction in expression of *FLC* on chromosomes 13 and 20 during cold treatment, which would trigger flowering, since *FLC* would be expected to suppress floral initiation. *FLC* on chromosome 8 also showed reduced expression in the *C. sativa* ssp. *pilosa* winter parent upon cold treatment, but was expressed at very high levels across all time points in the spring-type *C. sativa*. The chromosome 8 copy carried a deletion in the spring-type line, which could impact its functionality. Contrary to previous reports, all three *FLC* loci can contribute to controlling the vernalization response in *C. sativa* and provide opportunities for manipulating this requirement in the crop.

**Significance Statement:** Developing winter *C. sativa* germplasm is an important breeding goal for this alternative oilseed, with application in the food, fuel and bioproduct industries. Studying the genetic architecture of the vernalization response has shown that contrary to previous reports all three *FLC* loci in *Camelina* species could be exploited to manipulate this important trait.

## Introduction

The evolutionary path to form *C. sativa* is believed to have created a genetic bottleneck leading to low genetic diversity in spring-type *Camelina* germplasm, (Vollmann *et al*., 2005, Gehringer *et al*., 2006, Singh *et al*., 2015, Luo *et al*., 2019a) that has hindered the efforts to improve Camelina through breeding. In addition, hybridization of this crop with other species has had limited success (Narasimhulu *et al*., 1994, Hansen, 1998, Jiang *et al*., 2009, Séguin-Swartz *et al*., 2013, Julié-Galau *et al*., 2014, Martin *et al*., 2015). Although, interspecific hybridization was successful between *C. sativa* and *C. microcarpa* and produced plants of intermediate phenology, the hybrids displayed low levels of pollen viability and reduced fitness (Martin *et al*., 2019). The use of wide crosses can be an important tool to increase the genetic diversity in a crop, as well as to identify potentially important quantitative trait loci (QTL). However, challenges to this approach can exist due to several factors, such as asynchronized flowering behaviour, fertility issues and fundamental differences in the chromosome number between species (Chaudhary *et al*., 2020). Identification of wild relatives (Martin *et al*., 2017, Brock *et al*., 2018), which are closely related to the domesticated *C. sativa* have encouraged their use in *C. sativa* breeding and the extent of relatedness among the *Camelina* species almost certainly plays a role in the success of hybridization. As might be expected, *C. sativa* sub-species, such as *C. sativa* ssp. *pilosa* (DC.) N.W. Zinger, and the closely related *C. alyssum* (Mill.) Thell. (also suggested to be a sub-species or even a synonym of *C. sativa*) show higher success in hybridization attempts relative to wild relatives, such as *C. microcarpa* (Séguin-Swartz *et al*., 2013, Martin *et al*., 2019).

Plants with winter growth habit invariably require vernalization, that is exposure to a period of low but non-freezing temperatures, to transition from the vegetative stage to the reproductive stage. Most *C. sativa* germplasm behaves as an annual, but among close relatives a few, including *C. sativa* ssp. *pilosa* and *C. alyssum*, have been characterized with a biennial growth habit (Galasso *et al*., 2015), yet they share the same number of chromosomes with hexaploid *C. sativa* (Chaudhary *et al*., 2020). A recent study of winter- and spring-types of *C. sativa* compared leaf morphology, growth behaviour and seed characteristics (Wittenberg *et al*., 2019), where marked reduction in leaf number, plant height and plant growth before vernalization were reported for winter-types. Winter-type *C. sativa* is hardy to adverse winter conditions and displays good stand establishment (Gesch *et al*., 2018), and is considered a suitable candidate for double- and relay-cropping on the Northern Great Plains (Berti *et al*., 2017). Vollmann and Eynck (2015) noted that changed oil composition with higher linolenic acid levels, early flowering and avoidance of a number of biotic and abiotic factors were some of the advantages associated with winter-type *C. sativa.* Also, a lower level of erucic acid, an anti-nutritional compound, has been reported in winter-type *C. sativa* compared to spring-type *C. sativa* (Kurasiak-Popowska *et al*., 2020). Thus far, there has been limited exploration of winter-type *C. sativa* germplasm that can survive prolonged harsh winters with similar yields as current spring-types.

A vernalization requirement is one of the major characteristics of winter-type plants and *Flowering Locus C* (*FLC*) has been identified as a major regulatory gene in the vernalization pathway (Michaels and Amasino, 1999, Swiezewski *et al*., 2009). FLC is responsible for supressing bolting in the plant, with a higher level of expression maintained in the winter-type that gradually reduces with duration of cold treatment (Anderson *et al*., 2018, Schiessl *et al*., 2019, Takada *et al*., 2019). The neopolyploid *C. sativa* possess three copies of *FLC* (Kagale *et al*., 2014). The potential for sub-functionalization of *FLC* in *C. sativa* cannot be ignored since it has been reported for other Brassicaceae polyploid species (Schiessl et al., 2019). The expression pattern of *FLC* on chromosome 20 (Csa20g15400) implied it has a function in differentiating winter-type and spring-type *C. sativa* (Anderson *et al*., 2018, Chao *et al*., 2019). Although Anderson et al (2018) suggested the additional copies of *FLC* might play alternative roles in *C. sativa,* more recently a quantitative trait locus (QTL) found on chromosome 8, encompassing the *FLC* region was shown to have an effect on flowering time in spring *C. sativa* lines (Li *et al*., 2021, Lily *et al*., 2021).

Various methods have been developed to detect QTL associated with a particular trait,among them, genome-wide association analyses (GWAS) has become popular to capture variation present in diverse populations. However, it can be difficult to manage the requisite large populations, in particular, phenotyping can be cumbersome. The development of biparental populations to identify QTL is an establised approach where only prior knowledge for a quantitative difference in a trait of interest among parents is required. With advancements in sequencing technologies, the time and cost associated with marker generation has been reduced (Hall, 2013). Likewise, availability of the *C. sativa* reference genome (Kagale *et al*., 2014) offers the potential to identify candidate genes controlling traits of interest (King *et al*., 2019, Luo *et al*., 2019b). In this context, genotyping-by-sequencing (GBS) is a valuable technique to generate genetic information at low cost (Poland *et al*., 2012) and can be used to create genetic linkage maps and map loci controlling traits of interest (Young and Tanksley 1989).

In this study, one spring-type *C. sativa* genotype was crossed with two different winter biotypes of *Camelina* (*C. sativa* ssp. *pilosa* and *C. alyssum*) to study the genetic mechanisms underlying vernalization requirement in winter-type *C. sativa.* All three *FLC* orthologs were identified as potential candidate genes controlling flowering in *Camelina* species. The original hypothesis was that the same QTL would control the vernalization requirement, irrespective of source; however, the results suggested that dependent upon the source of the winter phenotype, combinations of QTL originating from different subgenomes of the hexaploid act to determine the vernalization requirement in *C. sativa*.

## Results

### Population development and determination of winter-type behaviour in *Camelina*

The cross between spring-type *C. sativa* (TMP23992) and winter-type *C. sativa* ssp. *pilosa* (CN113692) produced a semi-winter hybrid which took 54 days to flower without vernalization in comparison to 30 days for the maternal *C. sativa* (TMP23992) and 87-91 days for the paternal *C. sativa* ssp. *pilosa* (CN113692). Crosses between *C. sativa* (TMP23992) and winter-type *C. alyssum* (CAM176) produced winter-type hybrid plants that similar to CAM176 required vernalization in order to flower.

Two F_2_ populations (Csp and Csa) were developed from the F_1_ hybrids (single hybrid plant for each population) derived from each cross and were used to determine the segregation of winter-type behaviour (**Figure S1**). For both populations, segregation of winter-type habit (based on leaf morphology and early plant growth) was noted in the F_2_ and their progeny F_2:3_ lines. Segregation for days to flower and reduced stem growth was observed for both populations.

In the case of the Csp population, 45 of 118 F_2_ lines showed spring-type behaviour, while the remaining 73 lines showed semi-winter-type behaviour (**Figure 1A**), suggesting a single dominant gene controlled flowering. The semi-winter lines were subjected to vernalization and all F_2_ lines flowered within 70 days of seeding, including a vernalization period of 15 days. A total of 96 F_2:3_ lines were grown and phenotyped from the Csp population, where all lines flowered within a range of 27-55 days of seeding without vernalization. There were no typical winter-type plants among the F_2:3_ lines; however, the lines segregated for days to first flower (**Figure 1C**).

**Figure 1.**
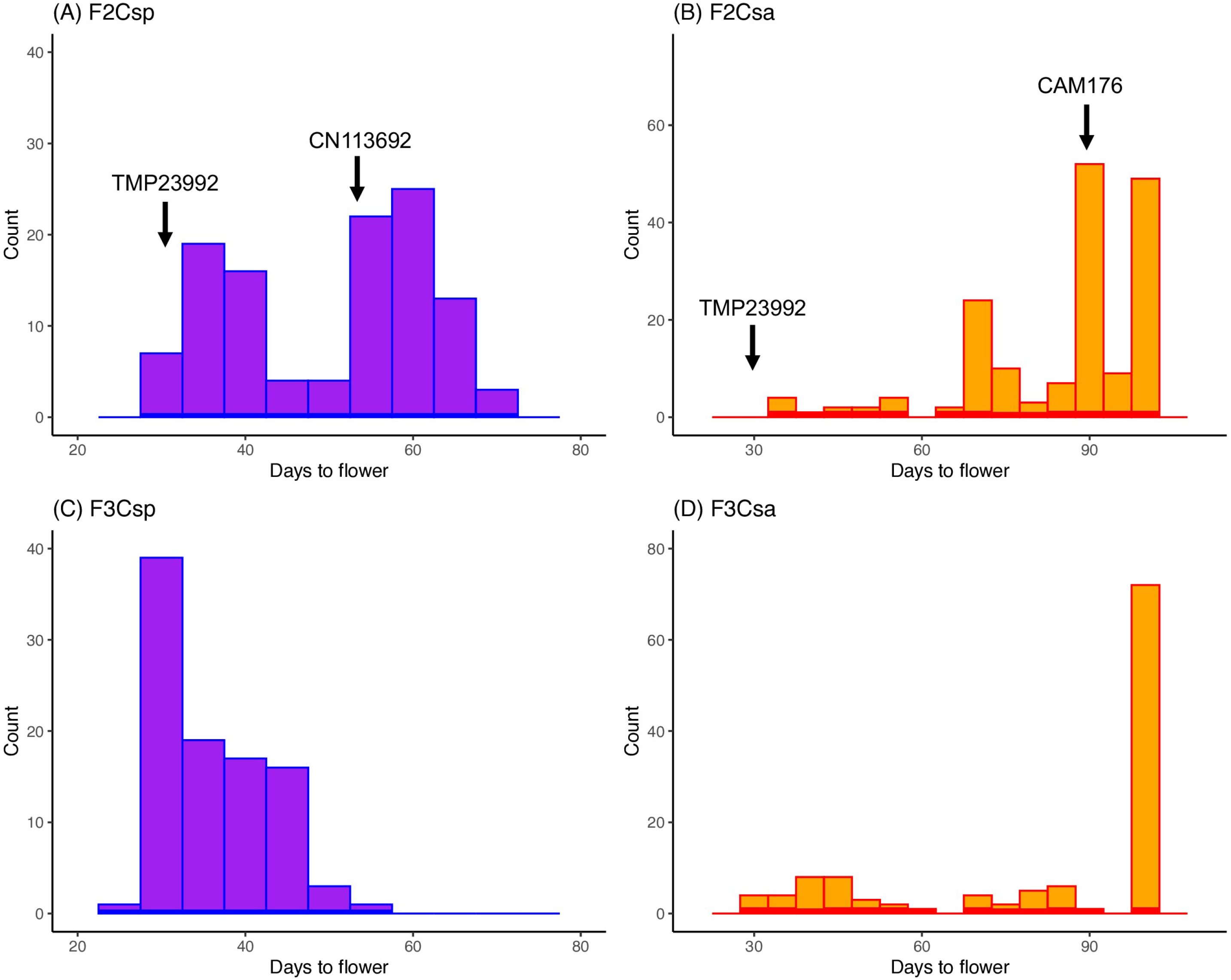
Flowering behaviour in segregating intra-specific *Camelina* populations. Frequency distribution of days to flowering in (A) F_2_ developed from a *C. sativa × C. sativa* ssp. *pilosa* cross (Csp); (B) F_2_ developed from a *C. sativa × C. alyssum* cross (Csa); (C) F_2:3_ developed from a *C. sativa × C. sativa* ssp. *pilosa* cross; and (D) F_2:3_ developed from a *C. sativa × C. alyssum* cross.

For the Csa population, 169 F_2_ lines were grown, of which 13 lines showed typical spring-type growth behaviour and the remaining 156 lines showed winter-type behaviour (based on reduced stem elongation) (**Figure 1B**), suggesting multiple loci controlled flowering. All 156 lines were subjected to vernalization treatment for 30 days; however, only 126 lines flowered within 100 days of seeding. From these F_2_ lines, 120 were successfully established in the F_2:3_ generation, which were tested for flowering behaviour in the absence of vernalization treatment. Among the 120 F_2:3_ lines, 30 lines were identified as spring-type and produced flowers within 58 days of seeding, 18 lines transitioned to the flowering stage with only a few flowers after 70 days of seeding, while 72 lines did not produce any flowers until at least 100 days after seeding (**Figure 1D**).

### Genetic linkage maps of *Camelina sativa*

For genotyping, 118 F_2_ lines from the Csp population and 169 F_2_ lines from the Csa population were used. In the case of the Csp population, 84,346 SNPs were identified and after filtering for those with more than 10% missing genotypes or distorted segregation ratios; 1,550 SNPs were used to construct a genetic linkage map (**Figure 2A**). Although attempts to include additional SNP was made by increasing the threshold for missing genotypes, this led to significant deviations from the expected segregation ratio, which could suggest errors in the genotype calls. A linkage map with a total length of 2193.8 cM was constructed, where the number of markers per linkage group ranged from 16 on chromosome 2 to 158 on chromosome 20 with an average mapping interval of 1 marker per 1.42 cM (**Table S1**). For the Csa population, 115,827 SNPs were identified for 169 genotypes; however, only 96 genotypes with sufficient sequence coverage to confidently call SNPs were used to generate the genetic linkage map. Upon filtering for distorted segregation and those with more than 5% missing genotypes, 3,279 SNPs were identified and mapped across the 20 chromosomes of the reference *C. sativa* genome (**Figure 2B**). The map encompassed 2399.96 cM with an average of 0.73 cM/marker. The number of markers per linkage group ranged from 50 on chromosome 2 to 420 on chromosome 11 (**Table S2**).

**Figure 2.**
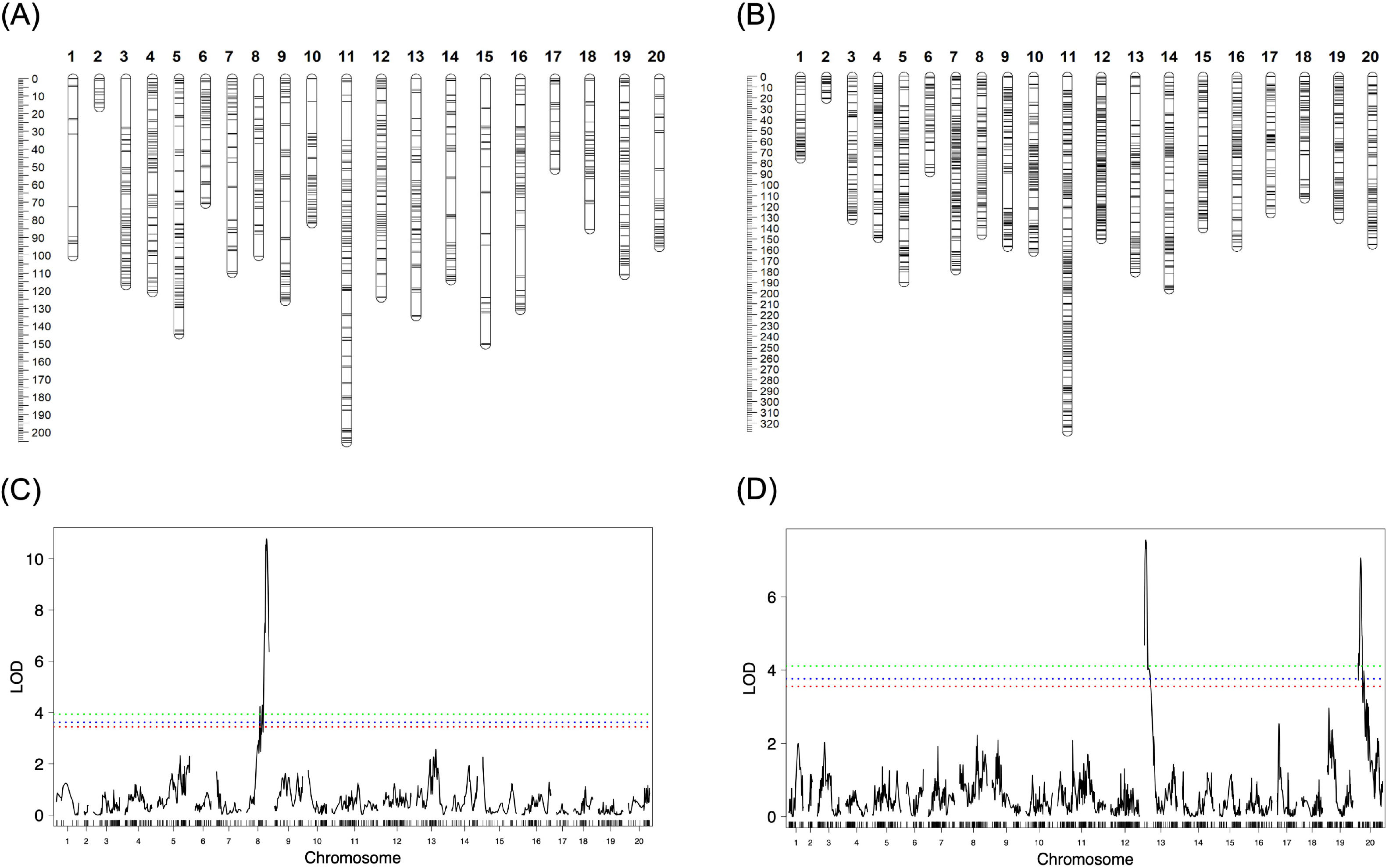
Mapping of QTL associated with vernalization requirement in C. sativa. (A) Genetic linkage map derived from the *C. sativa × C. sativa* ssp. *pilosa* (Csp) F_2_ population; (B) Genetic linkage map derived from the *C. sativa × C. alyssum* (Csa) F_2_ population; (C) QTL identified in the *C. sativa × C. sativa* ssp. *pilosa* (Csp) F_2_ population; and (D) QTL identified in the *C. sativa × C. alyssum* (Csa) F_2_ population. cM distance is shown to the left of the maps in panels (A) and (B). The significance threshold for identifying QTL is shown as a green line in panels (C) and (D).

The two genetic maps showed good collinearity along their length (**Figure S2**). In addition, the genetic maps identified a potential miss-assembly in the reference genome of *C. sativa* on chromosome 16 (∼10 Mb region), where an insertion from the terminal region of chromosome 17 (34 Mb) was found for both maps. The inserted region represented a small fraction of ancestral genomic block D (Kagale *et al*., 2014).

### Mapping QTL for winter-type behaviour in *Camelina*

For both populations QTL were identified using days to flower (DTF) values measured for the F_2_ lines, where the data represented variation in DTF in response to vernalization. For the Csp population, the analysis identified a strong QTL correlated with winter-type behaviour on chromosome 8, base pair position 2,323,768 (LOD = 10.8), which explained 36.07% of the phenotypic variation (**Figure 2C**) (**Table 1**). The *C. sativa* ssp. *pilosa* allele was co-dominant, where heterozygosity at the linked SNP loci was associated with an intermediate late flowering phenotype (**Figure S3**). The range of the confidence interval for the identified QTL was 7 cM. In the physically mapped region (0.18 Mb), within the 95% confidence interval of the QTL, 562 annotated genes were identified. An ortholog of *FLC* (*Csa08g054450*) was identified, which was just 65 kb away from the QTL peak and 15 further flowering-related genes were identified within or close to the QTL region (**Table S3A**).

**Table 1.**
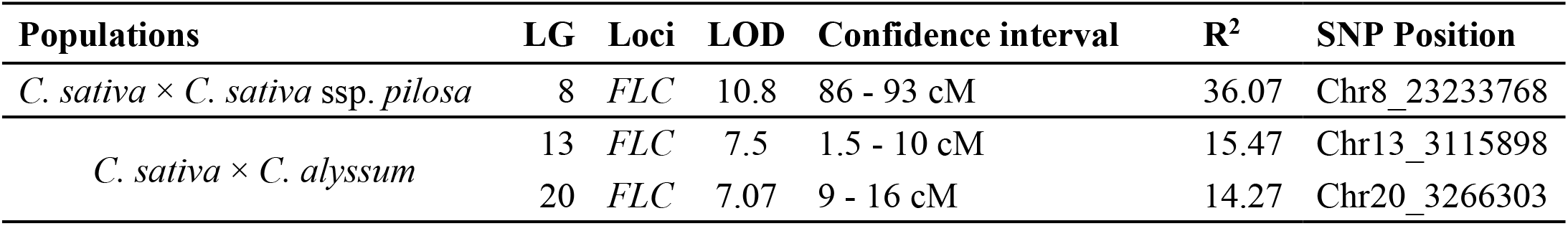
QTL for vernalization requirement in C. sativa measured as a days to first flower in F2 populations.

| **Populations** | **LG** | **Loci** | **LOD** | **Confidence interval** | **R<sup>2</sup>** | **SNP Position** |
| --- | --- | --- | --- | --- | --- | --- |
| *C. sativa* × *C. sativa* ssp. *pilosa* | 8 | *FLC* | 10.8 | 86 - 93 cM | 36.07 | Chr8_23233768 |
| *C. sativa* × *C. alyssum* | 13 | *FLC* | 7.5 | 1.5 - 10 cM | 15.47 | Chr13_3115898 |
|  | 20 | *FLC* | 7.07 | 9 - 16 cM | 14.27 | Chr20_3266303 |

Two QTL were identified in the Csa population, one on chromosome 13 (LOD = 7.50) and the second on chromosome 20 (LOD = 7.07) (**Figure 2D**) (**Table 1**). The QTL map interval on chromosome 13 was 8.5 cM (3.07 Mb on the physical map), whereas it was 7 cM (3.3 Mb) on chromosome 20. In this population, the QTL on chromosome 13 showed a dominant effect, whereas that on chromosome 20 showed a co-dominant effect (**Figure S3**). These two QTL intervals represented homoeologous segments of the reference *C. sativa* genome, where the QTL interval on chromosome 13 comprised 717 genes and that on chromosome 20 comprised 630 annotated genes. The QTL interval on chromosome 13 represented the terminal region of the linkage group and encompassed 20 flowering related genes (**Table S3A**). Beside this, the peak of the QTL was 913 Kb away from an ortholog of *FLC* (*Csa13g011890*). In the case of the QTL on chromosome 20, 17 genes were identified as flowering-related genes within the confidence interval of the QTL (**Table S3A**), among these *FLC* (*Csa20g015400*), *EMF1* (*Csa20g017070*) and *FY* (*Csa20g018850*) were identified to have a role in the vernalization response (Michaels and Amasino, 1999, Aubert *et al*., 2001, Simpson *et al*., 2003). These two QTLs together explained 29.74% of the phenotypic variation in the Csa population (**Table 1**).

All three identified QTL loci were found in homoeologous segments of the reference *C. sativa* genome, showing synteny with *A. thaliana* chromosome 5 ancestral genome block R, but with a slight difference in the absolute position of the QTL confidence interval (**Figure S4**). The one gene found in proximity to all three QTL with a defined role in the plant’s vernalization response was *FLC,* suggesting that it could be the probable candidate gene for this requirement in *C. sativa*.

### Differential gene expression in *Camelina* at the QTL loci

Gene expression analysis was performed at different time points within the parental lines: *C. sativa* (TMP23992) spring-type, *C. sativa* ssp. *pilosa* (CN113692) winter type and *C. alyssum* (CAM176) winter-type. For spring-type *C. sativa,* plants were kept in vernalization for one week, longer proved detrimental to plant development; plants were sampled after one week in vernalization (1W). For the two winter-types (*C. alyssum* and *C. sativa* ssp. *pilosa*), plants were kept in vernalization for four weeks, with tissue sampling at two time points (two weeks (2W) and 4 weeks in vernalization (4W)). Gene expression levels were compared before-vernalization (BV) versus 1W and 1W versus one week post-vernalization (PV) for spring type *C. sativa*. Likewise, comparisons were made for the two winter-types at BV versus 2W, 2W versus 4W, and 4W versus PV. The significantly differentially expressed genes were compared in the identified QTL regions on chromosomes 8, 13, and 20 to identify genes distinguishing the winter- and spring-type parents.

Gene expression analysis across the three time points in spring-type *C. sativa* identified 9,224 differentially expressed genes (*P*-value<0.05), among these 186 were related to genes associated with flowering time in other species (**Table S3B**). In the case of *C. alyssum*, 378 genes were differentially expressed across all time points, among these 14 genes were related to flowering; and in *C. sativa* ssp*. pilosa* 1,544 genes were differentially expressed across all time points, where 67 genes were associated with the flowering response (**Figure 3**, **Table S3B**). In the QTL region on chromosome 8, of those showing average expression levels of >5 FPKM in at least one sample, 3 genes showed significant differential gene expression across different time points in *C. sativa* ssp. *pilosa* **(Table S4**). *FLC* was the only flowering related gene found in the QTL region and was down-regulated in response to the vernalization treatment with expression remaining low post-vernalization, unlike in the spring-type parent where expression of *FLC* was high post-vernalization. Interestingly, *C. alyssum* showed the same expression pattern for the *Csa.FLC.C08* (*Csa08g054450*) copy. In the QTL regions on chromosome 13 and chromosome 20, 28 and 20 genes, respectively were found to be differentially expressed at any one time point in *C. alyssum* (**Table S5)**. Among these only one gene from chromosome 13 and three genes from chromosome 20 showed differential gene expression across all time points (**Figure 4, Table S5**). In both instances *FLC* was found to be down regulated in response to vernalization. In the case of spring-type *C. sativa*, the genes showing differential expression in the QTL regions of the winter and semi-winter type were not similarly expressed, notably although *Csa.FLC.C08* showed a slight reduction in expression after one week of cold treatment, expression increased post-vernalization (**Table S6**).

**Figure 3.**
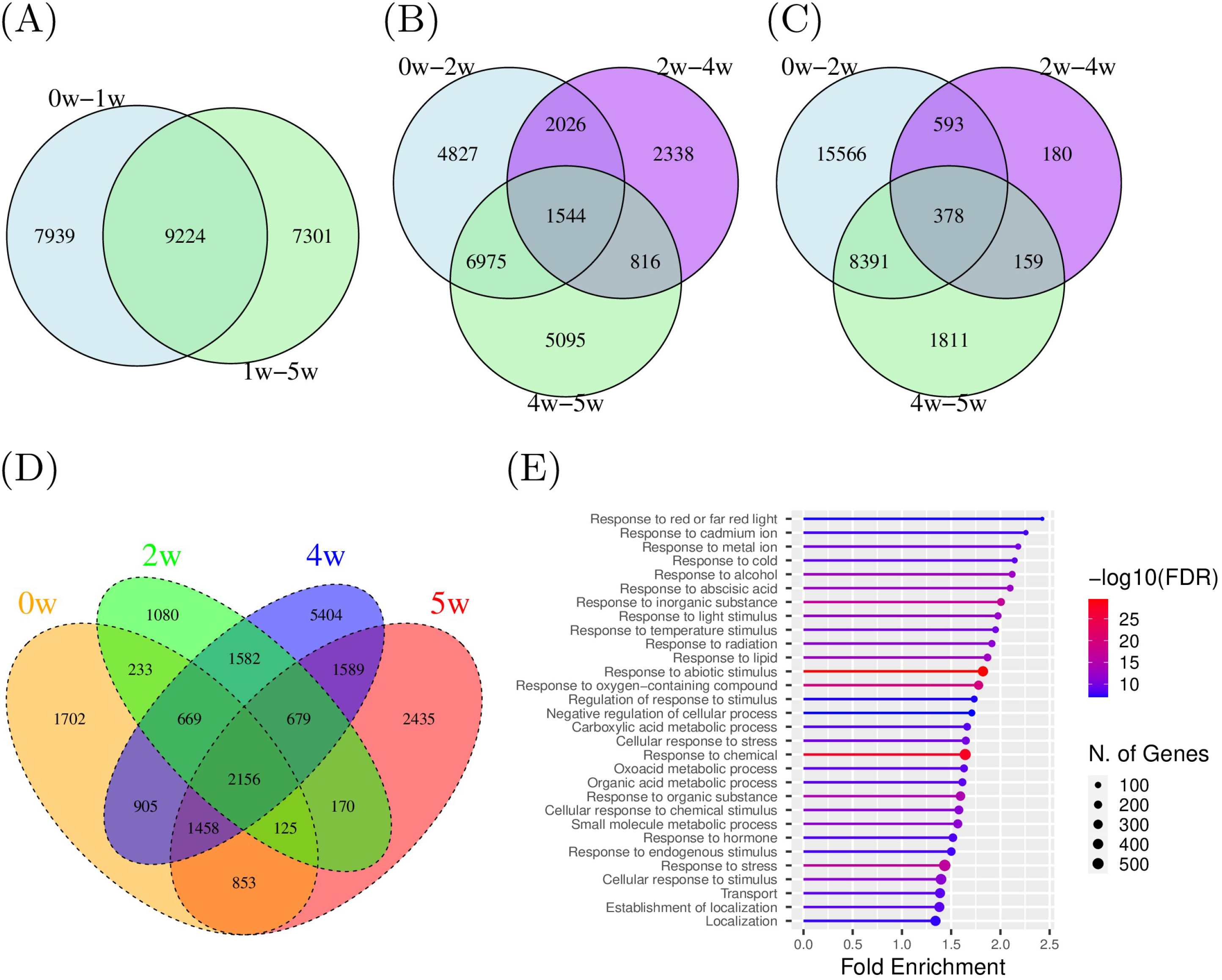
Differential gene expression before, during and after vernalization in spring- and winter-type *Camelina* parents. Genotype TMP23992 (*C. sativa*) is a spring-type, whereas CAM176 (*C. alyssum*) and CN113692 (*C. sativa* ssp. *pilosa*) are winter-types. Venn diagrams showing number of differentially expressed genes (DEGs) across time points in (A) TMP23992, (B) CN113692, and (C) CAM176; where BV is before-vernalization, 1W, 2W, and 4W are one, two or four weeks during vernalization, and PV is one week post-vernalization. Venn diagram showing number of DEGs between two winter-type *Camelina* genotypes, CN113692 and CAM176, at 4 different time points (D). Gene ontology analysis of unique genes expressed after four weeks of vernalization in CAM176 (E).

**Figure 4.**
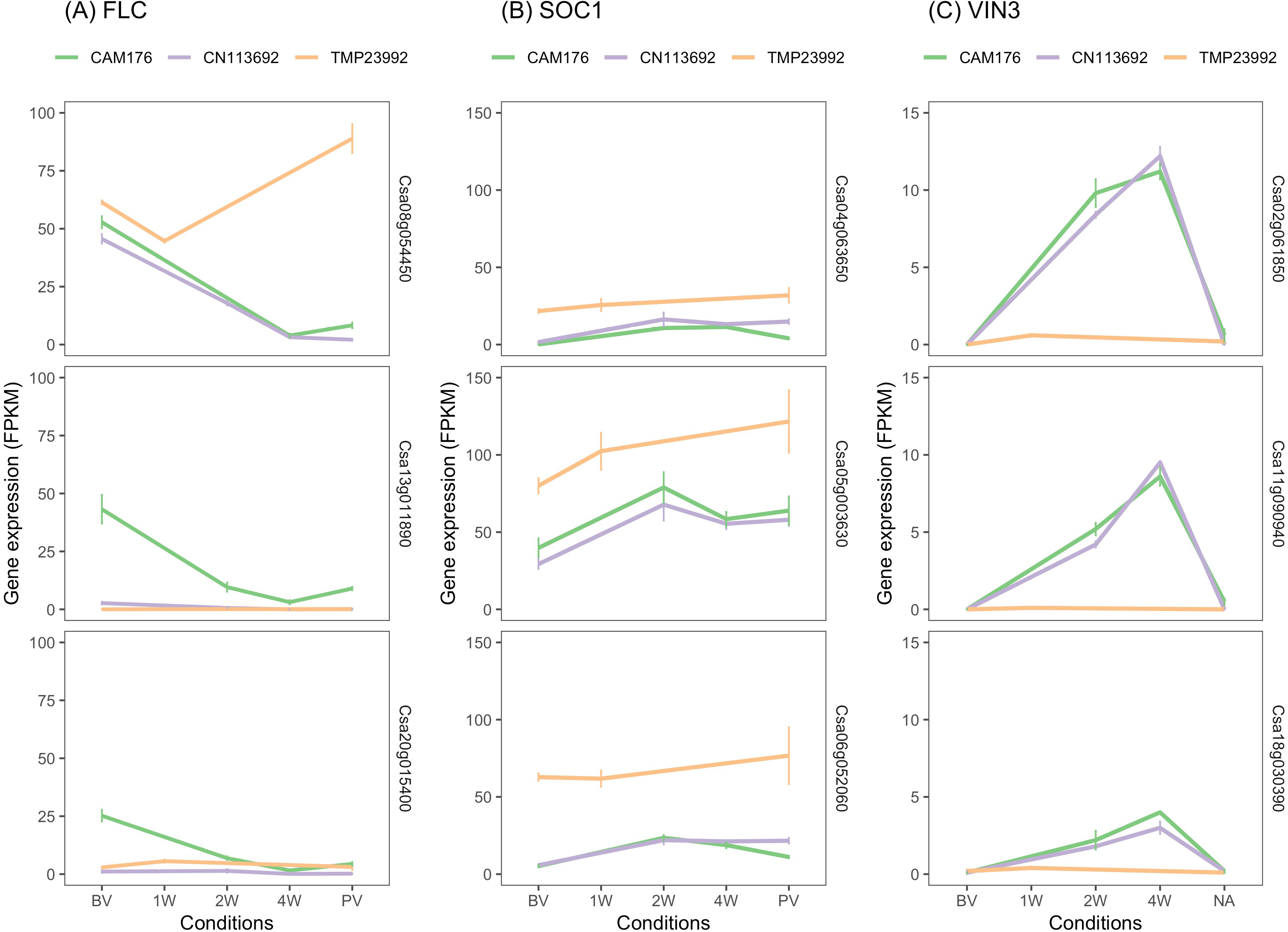
Changes in the gene expression during different time points of vernalization in Camelina species. Gene expression levels (FPKM) of each ortholog of *FLC* (A), *SOC1* (B), and *VIN3* (C) in three parental lines. FPKM represents mean Fragments Per Kilobase of transcript per Million mapped reads calculated for replicated RNASeq data; error bars represent standard error of the mean value.

Since the two winter type lines showed differing phenotypes with *C. sativa* ssp. *pilosa* being a semi-winter-type compared to *C. alyssum,* a true winter type, the two winter-types were compared at each time point; 2,156 genes were found to be differentially expressed across all time points (**Figure 3D**). Notably, the level of differential gene expression between the two genotypes increased with duration of vernalization, with 1,080 and 5,404 genes uniquely differentially expressed after two and four weeks of vernalization, respectively (**Figure 3D**). The gene ontology of these genes suggested the majority are related to response to stimuli and stress due to changes in the environment (**Figure 3E**). This might indicate a varying level of response to cold treatment in the two genotypes.

Gene co-expression analysis using weighted gene co-expression network analysis (WGCNA) was performed to identify genes that were clustered together with the major flowering gene *FLC*. In the case of the spring-type *C. sativa* (**Figure S5A**), *FLC* was absent from all 11 modules identified (**Table S7A**). For *C. sativa* ssp. *pilosa FLC* from chromosomes 8 and 20 clustered in one group along with 242 other genes (**Table 7B, Figure S5B**). Likewise, in *C. alyssum FLC* from chromosome 8, chromosome 13, and chromosome 20 clustered together along with 32 other genes (**Table S7C, Figure S5C)**. However, comparing the modules containing *FLC* between the two genotypes identified only 8 genes in common, suggesting divergence in the co-expression of related genes between these two winter-type *Camelina* species.

### Pathway for flowering in *Camelina*

It has previously been shown that *C. sativa* and its diploid progenitor *C. neglecta* do not appear to contain an ortholog of a key gene in the *A. thaliana* flowering pathway, *FRIGIDA* (*FRI*) (Chaudhary et al, 2022). Thus the expression of flowering-related genes was explored to infer whether the well-defined pathways of *A. thaliana* (Teotia and Tang, 2015) were in fact good predictors for gene expression in *C. sativa* (**Figure 5; Table S8A**). Only expressed genes (FPKM>0.1 in at least two replicates) and those found in syntenic positions were considered in the final list (**Table S8B**). Thus 1,436 orthologs representing a comprehensive list of 579 *A. thaliana* genes known to be associated with flowering were identified in *C. sativa* and their expression profiles studied (**Table S8B**). Flowering is known to be controlled by a complex interplay of gene pathways, notably the photoperiod, the vernalization, the autonomous and the gibberellin (GA) pathways (Mouradov *et al*., 2002). Adding further complexity, expression of most flowering time genes are also impacted by the circadian rhythm pathway, being differentially expressed between day and night (Fowler *et al*., 1999, Mizoguchi *et al*., 2005). Inferring function from expression is limiting but it was apparent that orthologues of almost all known *A. thaliana* flowering time genes were identified among the differentially expressed genes. Of note, in addition to *FRI*, a further well characterised flowering time gene, *Flowering Wageningen* (*FWA*), was also completely absent from the *C. sativa* genome. Clustering of the significantly differentially expressed key flowering related genes identified three clusters (**Figure 5A**). The first cluster shows reduced expression during vernalization while the second shows the opposite response. The latter cluster also showed variation in expression across biotypes as well as with the time period of cold treatment, with three sub-patterns within the cluster; those more highly expressed in the spring-type, those expressed predominantly in the two winter-types, and those expressed transiently in the winter types. The third small cluster contained genes which were largely repressed during vernalization and differentiated among the biotypes. Cluster 1 was largely defined by those genes with higher expression post-vernalization in the spring type. The expression patterns clearly differentiated the spring- and winter-types with some notable genes highlighted in **Figure 5B**. However, except for expression of two copies of *FLC*, those on chromosome 13 and 20, differences between the winter-types was more subtle. In the *C. alyssum* line, some of the photoperiod and circadian genes maintained higher levels of expression throughout the cold treatment compared to the semi-winter line (eg. *CCA* and *COL1* orthologs). As perhaps expected from the undifferentiated nature of the *C. sativa* genome, most *A. thaliana* flowering-time genes were maintained in three copies, interestingly there was limited genome divergence of gene expression pattern or genome dominance observed among the duplicate copies (**Figure S6**).

**Figure 5.**
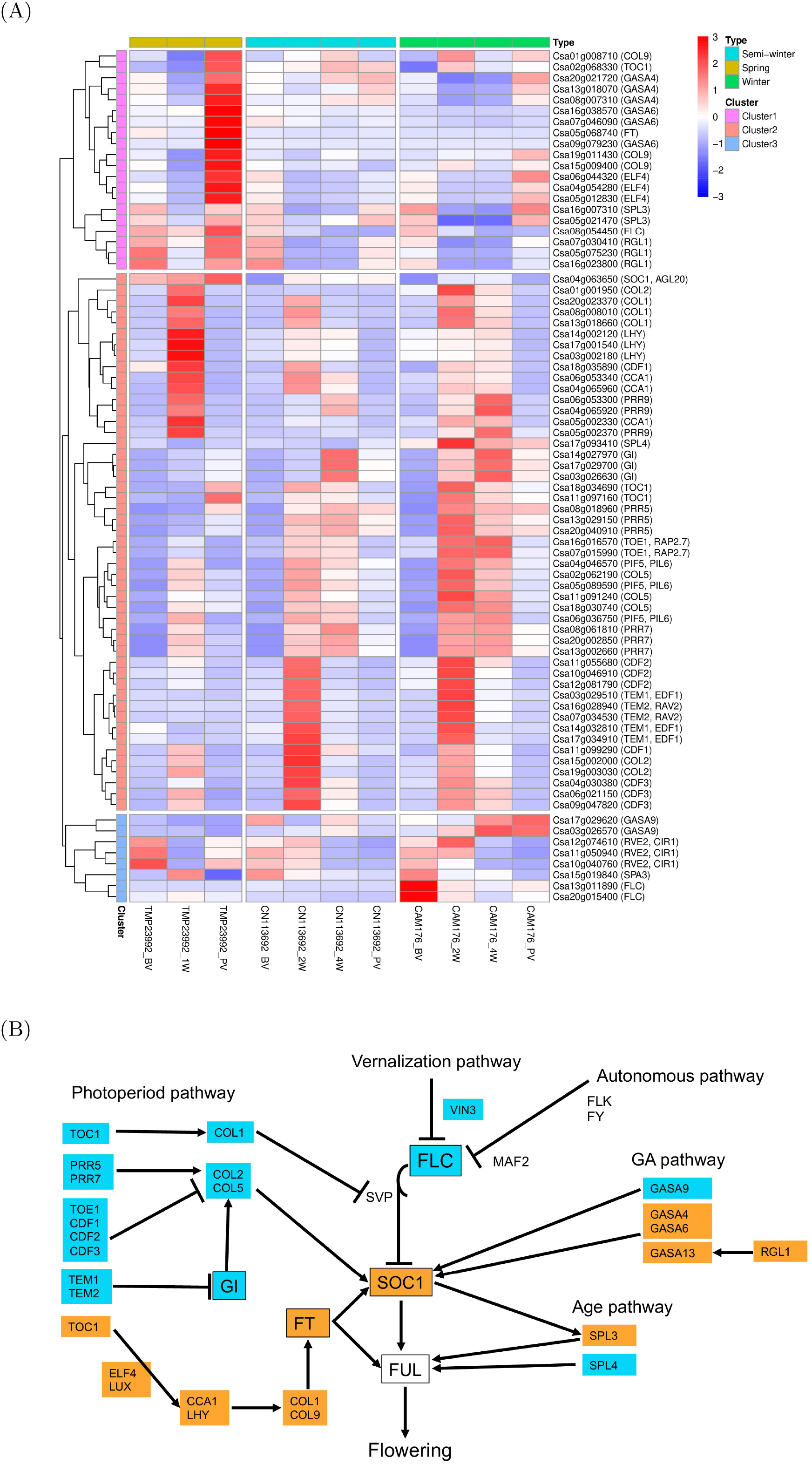
Flowering pathway in Camelina sativa in response to vernalization treatment. A) Heatmap showing expression of flowering related genes across predefined flowering pathways in *A. thaliana* (Photoperiod, Vernalization, Autonomous and Gibberellin) and in *Camelina* genotypes TMP23992 (spring), CN119243 (semi-winter) and CAM176 (winter) for different level of cold treatment (BV- pre-vernalization, 1W- 7 days at vernalization, 2W, 14 days at vernalization, 4W- 30 days at vernalization, and PV- 7 days post vernalization); and B) A schematic diagram for flowering pathway in *C. sativa* where flowering related gene showing differential gene expression for at least one paralog in *C. sativa* are shown. The bar indicates negative effect, and the arrow indicates positive effect. Those genes highlighted in blue show differential expression in winter-type(s) while those in orange are differentially expressed in the spring type.

## Discussion

Flowering is a crucial stage in a plant’s growth cycle and has a direct influence on adaptation, fitness and overall plant productivity. In nature, biennial and annual flowering behaviour has been reported in a number of *Brassica* species and cereal crops (Kim *et al*., 2009). The annual nature of flowering is often characterized as an important adaptive trait during the domestication process in crop species (Ågren *et al*., 2017). The study of vernalization in *C. sativ*a could provide insights in other *Brassica* crops due to the high degree of homology shared among these species, as well as the close relationship of this species with the model plant *A. thaliana*.

Winter-type *Camelina* species are represented by plants requiring prolonged cold treatment to promote bolting. In this study, two different winter-type *Camelina* ecotypes were exploited. The first, *C. alyssum*, prior to vernalisation had shorter stems characterized by profuse leaf production, where cold treatment promoted stem elongation and flowering. The second type, *C. sativa* ssp. *pilosa*, was charaterized by longer stems and branching and might be considered a winter annual, since flowering would occur without vernalization, but was induced more rapidly and profusely after 2-3 weeks of vernalization (**Figure S7**). The availability of these two forms of winter-type *Camelina* enabled potentially different mechanisms controlling delayed flowering in *C. sativa* to be studied. Hybrids generated between *C. sativa* and *C. alyssum* produced an obligate winter-type plant, suggesting a dominant trait. A number of reports have shown a quantitative effect for duration of vernalization (Sheldon *et al*., 2000, Kemi *et al*., 2013). Similar observations were made in this study, as the days to flowering was greater for hybrid plants (*C. sativa × C. alyssum*) vernalized for a shorter duration compared to those vernalized for a longer period (**Figure S8**). The extent of variation in vernalization requirement, as reflected by days to flowering, as well as the difference in the number of reproductive branches in hybrids coming from the same parents (**Table S9**), suggested there was quantitative variation for vernalization requirement.

Genetic linkage maps developed from populations derived from crosses between one spring- and two winter-type parents were aligned based on 648 common markers and showed a Spearman correlation coefficient of 0.81 (**Figure S2**), suggesting high contiguity of the maps. Since these populations were developed from crosses between one spring-type *C. sativa* parent with two different winter-type *Camelina* species/sub-species, the level of similarity shared among the winter-type parents influenced the number of common markers in the genetic maps. The maps both suggested mis-assembly of the reference genome, where a linkage block representing chromosome 17 was found on chromosome 16. A revised subgenome structure of the *C. sativa* genome had previously revealed that chromosomes 16 and 17 should be in different subgenomes (Chaudhary et al, 2020), and the resultant new syntelog table suggested genes belonging to the terminal region of chromosome 17 (subgenome 3) should be present in subgenome 2, which would be consistent with the results presented here (**Table S10**).

Three major QTL affecting winter-type behaviour in *C. sativa* were identified. The major QTL identified on chromosome 8 (subgenome 1) for the Csp population was in close proximity to an orthologue of *FLC,* indicating its potential role in flowering behaviour in *C. sativa* ssp. *pilosa.* Of note, although the F_2:3_ Csp population was not vernalized, all lines flowered within 55 days; yet the DTF phenotype data from the F_2:3_ lines identified the same QTL on chromosome 8 that controlled variation in DTF as reflected by the winter-type behaviour of the F_2_ lines (**Figure S9**). In contrast, two QTL were identified in the Csa population, where both QTL represented homologous regions in different subgenomes (chromosome 13: subgenome 2 and chromosome 20: subgenome 3). Within the confidence interval of these two QTL, or in close proximity, an orthologue of the major flowering time gene *FLC* was identified. However, low linkage disequilibrium (LD) detected for the markers around *FLC*, especially on chromosome 8, could suggest other genes might also be responsible for affecting days to flower (**Figure S10**). These QTL were confirmed through mapping of F_2:3_ phenotypes, where the same QTLs were identified on chromosomes 13 and 20 as in the F_2_ generation, but with a less significant *P-value* for the QTL on chromosome 20, which might suggest further segregation of codominant alelles (**Figure S11**). The low number of samples in both populations probably decreased the level of confidence for the identified QTL and in quantifying minor QTLs; however, the study identified three major QTL in two populations that have a significant effect in causing variation for flowering time/vernalization requirement.

A number of genes have been identified as being responsible for the vernalization requirement in *A. thaliana* and other related *Brassica* species. Among them, *FLC*, a well-characterized gene, has been shown to control winter-type behaviour in *A. thaliana* (Michaels and Amasino, 1999, Swiezewski *et al*., 2009). Orthologs of *FLC* have been shown in a number of *Brassica* species to affect vernalization requirement (Anderson *et al*., 2018, Schiessl *et al*., 2019, Takada *et al*., 2019), where higher expression of *FLC* suppresses flower initiation before vernalization. As such, the duration of vernalization is inversely correlated with the level of *FLC* expression over the course of the vernalization period (Sheldon *et al*., 2000), and *FLC* acts as a repressor for a number of genes associated with flowering responses (Deng *et al*., 2011). *Camelina sativa* is a hexaploid with three relatively undifferentiated subgenomes which is reflected by the existence of three copies of *FLC* (Kagale *et al*., 2014). Previously, one ortholog of *Csa*.*FLC.C20* (*Csa20g015400*) was found to be differentially expressed in response to vernalization in the winter-type *C. sativa* variety Joelle in comparison to spring-type *C. sativa* (Anderson *et al*., 2018). This was confirmed by Chao *et al*. (2019) with an additional set of winter-type *C. sativa* lines, where expression of *Csa*.*FLC.C20* could differentiate the two biotypes. It was speculated that the additional *FLC* orthologs might have succumbed to selection pressure that resulted in a change or loss of function and they may now play a role in seed and/or tissue development (Anderson *et al*., 2018). Similarly, potential sub-functionalization of the *FLC* orthologs/homoeologs has been reported in some *Brassica* species (Schiessl *et al*., 2019). However, other studies have suggested that the additional *FLC* genes are responsible for variation in flowering time in the absence of vernalization requirement (Zou *et al*., 2012, Xiao *et al*., 2013, O’Neill *et al*., 2019)

The proximity of *Csa.FLC* to the identfied QTL suggested that *Csa.FLC* could be influencing vernalization requirement, as well as affecting days to flowering in winter-type *Camelina* similar to other crops (Okazaki *et al*., 2007, Zhao *et al*., 2010, Deng *et al*., 2011, Xiao *et al*., 2013). The QTL on chromosome 13 was an additional locus to those previously reported. In Chao *et al*. (2019), differential expression was observed for both *Csa.FLC.C08* and *Csa.FLC.C20*, while Anderson *et al*. (2018) suggested that *Csa.FLC.C20* was the determinant for vernalization requirement in *C. sativa*, where they identifed a one base deletion which resulted in a non-functional *FLC* protein in spring-type *C. sativa.* Reconstructing *Csa.FLC.C20* from the available RNAseq data indicated that the spring type TMP23992 shared homology with the published reference genome DH55 leading to a predicted truncated protein compared to the winter-type *C. alyssum* (**Figure S12**). Notably, *Csa.FLC.C08* in both the spring type TMP23992 and the reference genotype had a three base-pair deletion in comparison to both winter types, which was somewhat unexpectedly also reported in another winter-type *C. sativa* variety, Joelle (Anderson *et al*., 2018). The *FLC* ortholog in *Camelina neglecta*, a diploid progenitor of *C. sativa* aligns with the winter-type *Csa*.*FLC.C08* suggesting this to be the ancestral copy (Chaudhary *et al*., 2022) (**Figure S12**)*. Csa.FLC.C13* could only be reconstructed from *C. alyssum* and showed only synonymous sequence variation compared to the reference genome. For *C. alyssum*, the decrease in expression of *Csa.FLC.C13* and *Csa.FLC.C20* upon cold treatment suggested a similar role for *FLC* as that reported in a number of other species (Sheldon *et al*., 2000, Okazaki *et al*., 2007, Anderson *et al*., 2018). *Csa.FLC.C08* was also repressed in *C. sativa* ssp. *pilosa* as might be expected if it played a formative role in vernalization. *Csa.FLC.C08*, the likely progenitor copy, behaved as a functional copy in *C. alyssum*, with reduction in expression upon vernalization; although no significant QTL could be identified, there was a peak observed on chromosome C08 when studying the F_2:3_ population (**Figure S11**). Genome dominance has been observed in *C. sativa* with orthologs from the third sub-genome generally showing a higher level of expression (Chaudhary et. al, 2020), and this may be reflected with the capture of QTL in this instance. All three orthologs of *Csa.FLC* appear to have the potential to contribute to the control of flowering time in *C. sativa*, with each being differentially adapted in the spring-type, either through sequence variation or presumably control of gene expression in the case of *Csa.FLC.C13*. Interestingly, the variant copy of *Csa.FLC.C08* was found to be highly expressed in the spring type with expression increasing post-vernalization, suggesting it may have evolved a novel function in spring-type *C. sativa*.

The use of different winter-type sources helped to identify three homologous QTL responsible for winter-type behaviour in *C. sativa*, noteably, they were independently identified on different subgenomes in two populations. These loci through the generation of locus specific markers can be exploited in current efforts to develop winter-type *C. sativa* varieties, based on the different winter biotypes it appears that at least two of the identified loci might be required to develop true winter types. In conjunction with gene expression analyses, the three orthologs of *FLC* were identified as the likely candidates contributing to the variation underlying the QTL. Notwithstanding the absence of *FRIGIDA* (Chaudhary *et al*., 2022) and *FWA*, *C. sativa* was observed to share many of the key genes regulating both flowering time and the vernalization response with *A. thaliana* (**Figure 5**), further many of those genes maintained three orthologs in *C. sativa* and were similarly expressed (**Table S8**). As presented here, the presence of multiple orthologs can allow various routes for adaptation in the polyploid, thus further studies of flowering time in *C. sativa* may yet yield insights into the complexities of regulating this fundamental trait in new polyploid species.

## Experimental procedures

### Plant materials

Three different species were used to generate F_2_ and F_2:3_ populations: viz. *C. sativa* (TMP23992), *C. alyssum* (CAM176), and *C. sativa* ssp. *pilosa* (CN113692) (**Figure S1**). TMP23992 is a spring-type line, while the other two are winter-types. The spring-type TMP23992 produced flowers within 30 days of seeding. The two winter-type lines differed in morphology for winter behaviour, CN113692 was similar to the *C. sativa* spring-type in the early growth stages, but with increased vegetative branching and reduced height prior to cold treatment. CAM176 was characterized by a reduced stem with profuse leaves where the vernalization treatment promoted stem elongation, as well as branching and flowering (**Figure S7)**.

As shown in **Figure S1**, manual crossing was performed with unopened fully developed buds, where TMP23992 (spring-type) was the maternal parent and the winter-types were pollen donors. After pollination, flowers were covered with an isolation bag. Seeds from mature pods were harvested and planted. The hybrids between TMP23992 and CAM176 produced a winter-type plant; whereas those between TMP23992 and CN113692 produced semi-winter type plants, which flowered in the absence of vernalization; however, with a lower number of reproductive branches relative to the parental lines. Self-seed of each hybrid represented the F_2_ populations (**Figure S1**). F_2_ lines showing winter-type morphology were vernalized at 4 °C for 30 days for the *C. sativa* × *C. alyssum* cross (Csa) and 15 days for *C. sativa* × *C. sativa* ssp. *pilosa* cross (Csp). All experiments were carried out in the greenhouse in a soil-less potting mixture (Stringam, 1971) amended with controlled release fertilizer (15-9-12 Osmocote PLUS, Scotts Fertilizer Company, Marysville, OH, USA) with 16/8 hr of light/dark conditions. Vernalization requirement was determined based on the growth habit 20 days after seeding, where reduced stems with profuse leaves were characteristic of winter-type behaviour. Single seed descent was adopted to generate F_2:3_ lines for additional confirmation of growth habit. F_2:3_ plants were not subjected to vernalization; those lines either not flowering or late flowering with reduced flower numbers were assumed to have a winter habit. Days to first flower (DTF) for all the lines was recorded from the date of seeding. Plants not flowering 100 days after seeding were assigned a value of 100 for QTL mapping.

### Genotyping of segregating populations

Young leaf tissue was harvested from all lines and kept at -80 °C until DNA extraction. DNA extraction was performed using the CTAB method as described in (Chaudhary *et al*., 2020) and GBS library preparation was as described by Poland *et al*. (2012) using *PstI* and *MspI* for reduced genome representation. Paired-end 125 bp sequencing was performed with multiplexed libraries on a Hiseq platform (Illumina, San Diego, CA, US). The sequences were de-multiplexed followed by trimming of low quality bases and adapters using Trimmomatic version 0.33 (Bolger *et al*., 2014) where reads with a minimum length of 55 bp were retained. All high quality reads were mapped to the *C. sativa* reference genome (Kagale *et al*., 2014) using BWA (Li and Durbin, 2009) with *bwa-mem* tool with default parameters. From the aligned BAM files SNPs were called using the *UnifiedGenotyper* tool in GATK version 3.2-2 (McKenna *et al*., 2010) with default parameters.

### Genetic analyses of segregating populations

For both populations, all markers polymorphic between the parents were considered, apart from those showing distorted segregation, i.e. deviation from 1:2:1 (χ^2^ test, *P-value*< 0.05). Genetic linkage maps were prepared using MSTmap (Wu *et al*., 2007).

For the Csa population, SNPs for 96 F_2_ lines with less than 5% missing genotypes were used to construct a genetic linkage map, where logarithm of odds ratio (LOD) score of 7, mapping threshold of 1 and mapping distance threshold of 1 cM settings were used to determine the number of linkage groups. Markers that failed to cluster with their presumed linkage group (LG) of origin, based on alignment to the reference genome, were forced to cluster with said LG using the single LG function in MSTmap.

For the Csp population, SNPs for 118 F_2_ lines with less than 10% missing genotype data were used for map construction with a LOD score of 6, mapping threshold of 1 and mapping distance threshold of 1 cM. As before, further grouping of linkage groups was performed for those markers originating from the same physical chromosome, but separated by high genetic distances. The genetic maps were visualized using MapChart v2.32 (Voorrips, 2002). The genetic maps were compared for contiguity using the online version of genetic map comparator (Holtz *et al*., 2017).

### Identification of QTL

QTL analysis was performed with the R/qtl package (Broman *et al*., 2003) in R statistical software (R Core Team, 2021). A single QTL model developed with the Haley-Knott regression method was used to identify QTL. The significance threshold (LOD value) was determined using 1000 permutations and α=0.05, above which QTL were assumed to be significant. The fitqtl method with the drop one term method was adopted for identifying phenotypic variation explained by the QTL, where the method analyzes sub-models to fit the best model, and the percent variance explained for the QTL was calculated by the formula *h^2^* = 1-10^-2(/n)LOD^. The confidence interval of the QTL was identified using Bayesian Credible Interval in the R/qtl package and genes within the confidence interval of the QTL were identified from *C. sativa* annotated genes (Kagale *et al*., 2014). Homoeologous chromosomes with QTL were further visualized using KaryoploteR package in the R software (Gel and Serra, 2017).

### RNA sequencing and differential gene expression analysis

RNA sequencing of the parents, with three biological replications, for four different time points for *C. sativa* ssp. *pilosa* and *C. alyssum* and three time points for *C. sativa* were performed. The leaves tissues were harvested pre-vernalization (for all parents), 1 week into vernalization (TMP23992), 2 weeks into vernalization (CAM176 and CN113692), 4 weeks into vernalization (CAM176 and CN113692), and 1 week post vernalization (for all parents). Total RNA was extracted using a standard Rneasy Plant Qiagen kit as described by the manufacturer with on-column DNA digestion. RNA was quantified using a Qubit (Thermo Fisher Scientific Inc., Walthan, MA) and the quality determined using an RNA Nano labchip on a Bioanalyzer (Agilent Technologies, Santa Clara, USA). Paired-end RNAseq libraries were constructed using the Illumina Stranded mRNA Prep kit, with 500 ng of RNA for cDNA synthesis followed by RNA library preparation. The final library quality was checked using a TapeStation (Agilent Technologies, Santa Clara, USA). Sequencing was done on an Illumina NovaSeq 6000 platform (2 × 250 bp).

Sequence data were filtered for low quality reads, short reads and adapter contamination using Trimmomatic v.0.39 (Bolger et al., 2014). Leading and trailing bases with quality below 10, an average quality of base below 20 with a sliding window of 4, and reads shorter than 51 bp were removed. All trimmed reads were aligned with the annotated *C. sativa* reference genome (Kagale et al., 2014) using STAR v.2.7.9a (Dobin *et al*., 2013) using default parameters, except for *– alignIntronMax* set at 10000 and *–outFilterMismatchNmax* set at 4. *GeneCounts* in STAR provided read counts per annotated gene. The statistics on input data, filtration and mapping are presented in **Table S11**. Normalization of read counts was done using the Fragment Per Kilobase of transcripts per Million mapped reads (FPKM) method using RSEM v.1.3.3 (Li and Dewey, 2011). Differential gene expression analysis was performed with the DESeq2 package (Love *et al*., 2014) with “apeglm” function (Zhu *et al*., 2018) in R statistical software (R Core Team, 2021) where all genes were expected to be differentially expressed among comparisons based on *P-adjusted* value <0.05. Further, ggplot2 (Wickham, 2016) was used to plot the expression of genes across multiple time points for three parental genotypes and IGV v.2.11.1 (Thorvaldsdóttir *et al*., 2012) was used to visualize the alignment of candidate genes.

The mapped reads across the *FLC* orthologs were extracted using samtools v.1.13 (Li *et al*., 2009) and reassembled using Trinity v.2.13.2 (Grabherr *et al*., 2011) and protein models were predicted using TransDecoder v.5.5.0 (https://github.com/TransDecoder/TransDecoder). Multiple sequence alignment was performed using Clustal Omega in an online platform provided by EMBL-EBI (Madeira *et al*., 2022).

WGCNA package (Langfelder and Horvath, 2008) in R was used to infer gene co-expression/modules based on the transcriptional response with the change in condition such as pre-vernalization, time points during vernalization, and post-vernalization. Only 4471 highly differentially expressed genes (based on 95 quantile) were used in the analysis. Modules were identified based on hierarchical average linkage clustering with minModuleSize set to 30 and the co-expression measure to the power β=12. Gene Ontology and biological function of differentially expressed genes were analysed using Metascape (https://metascape.org) (Zhou *et al*., 2019).

A list of 579 *A. thaliana* flowering related genes were prepared by combining FLOR-ID genes (Bouché *et al*., 2016) with those from Sasaki *et al*. (2015). Orthologs in *C. sativa* were identified based on the syntelog table for *Camelina sativa* (Kagale *et al*., 2014). A total of 1436 flowering related genes in *Camelina sativa* were found to have an expression of >0.1 FPKM for at least two biological replications for any one time point (**Table S8**). Further, the top orthologs from these genotypes showing at least two-fold change in the expression during vernalization as well as having expression of >5 FPKM for at least two replicates were used for the preparation of heatmap using package pheatmap in R (R Core Team, 2021) (**Table S8**). The differentially expressed genes for each parental line were subjected to fit into KEGG pathway using ShinyGo 0.76.3 (Ge *et al*., 2019) to infer any potential differences in the pathway genes. The subgenome dominance analysis was performed with 425 flowering related genes having three copies across three subgenomes. The analysis of variance was performed with expression data (FPKM) to identify the subgenome having a dominant expression pattern; and the percentage of genes for a subgenome having dominant pattern were plotted using package “ggplot2” (Wickham, 2016) in R software.

## Acknowledgements

We would like to thank Lily Tang and Annette Zatylny for their support in the greenhouse and preliminary field experiments (data not shown). This work was funded in part by a grant from the Global Institute for Food Security in Saskatoon and in part by the Canadian Crop Genomics Initiative at Agriculture and Agri-Food Canada.

## Supplementary Information

**Figure S1.** Inter- and intraspecific hybridization scheme adopted in this study with total number of lines for segregating populations.

**Figure S2.** Collinearity between the Csa (*C. sativa* × *C. alyssum*) and Csp (*C. sativa* × *C. sativa* ssp. *pilosa*) genetic maps using The Genetic Map Comparator.

**Figure S3.** Effect of parental alleles on days to flower.

**Figure S4.** Structure of QTL regions for vernalization requirement in *C. sativa*.

**Figure S5.** Gene co-expression analysis using WGCNA.

**Figure S6.** Subgenome dominance analysis with flowering related genes in Camelina species.

**Figure S7.** Plant growth in two winter-type Camelina lines without vernalization.

**Figure S8.** Effect of duration of vernalization on flowering in hybrid developed from *C. sativa* × *C. alyssum* after 1 week (left) and three weeks (right) into vernalization.

**Figure S9.** QTL mapping for days to flower from F_2:3_ derived from Csp (*C. sativa* × *C. sativa* ssp. *pilosa*) population.

**Figure S10** Linkage disequilibrium heatmap showing R^2^ for markers across the QTL regions on chromosomes 8, 13 and 20. The markers labelled in blue are those flanking *FLC*.

**Figure S11.** QTL mapping of days to flowering from F_2:3_ derived from Csa (*C. sativa* × *C. alyssum*) population.

**Figure S12.** Reconstruction of *Flowering Locus C* (*FLC*) genes using Trinity from TMP23992 (*C. sativa*), DH55 (*C. sativa*), CN113692 (*C. sativa* ssp. *pilosa*), and CAM176 (*C. alyssum*) and comparison with *FLC* orthologs from DH55 (reference genome) and *C. neglecta* (reference genome).

**Table S1.** Distribution of markers on linkage map from *C. sativa* × *C. sativa* ssp. *pilosa* F2 populations.

**Table S2.** Distribution of markers on linkage map from *C. sativa* × *C. alyssum* F2 populations.

**Table S3.** Number of upregulated and downregulated genes in contrasting growth conditions in Camelina species.

**Table S4.** List of genes in the QTL region (chromosome 8) showing differential expression at different time points in genotype CN113692.

**Table S5.** List of genes in the QTL region showing differential expression at different time points in genotype CAM176.

**Table S6.** List of genes in the QTL region (chromosome 8, chromosome 13 and chromosome 20) showing differential expression at different time points in genotype TMP23992.

**Table S7.** Different modules identified with WGCNA.

**Table S8A.** Orthologues of A. thaliana flowering-time genes

**Table S8B**: List of orthologous flowering related genes with an expression >0.1 FPKM for at least two biological replications in *Camelina sativa*

**Table S8C.** Expression of flowering related genes and RNA expression log fold change at different time points.

**Table S9.** Days to first flowering for the hybrids developed from *C. sativa* × *C. alyssum* (TMP23992 × CAM176).

**Table S10.** Revised syntelog matrix adopted from Kagale et al. 2014 and revised based on Chaudhary et al. 2020 (The highlighted genes represent gene under the QTL).

**Table S11.** Statistics of RNA Seq data and mapping.

## Data Statement

The RNASeq data has been deposited at NCBI under SRA submission SUB12901385.

## Author Contributions

RC carried out the population development, phenotype assessment and genotyping of the lines. RC also carried out the RNASeq analyses with assistance from EEH. RC and IAPP designed the experiments and drafted the initial manuscript. CE assisted with phenotyping of the lines. AGS provided additional resources for data analyses. All authors contributed to editing the final manuscript.

## References

Ågren, J., Oakley, C.G., Lundemo, S. and Schemske, D.W. (2017) Adaptive divergence in flowering time among natural populations of *Arabidopsis thaliana*: Estimates of selection and QTL mapping. Evolution, 71, 550–564.

Anderson, J.V., Horvath, D.P., Doğramaci, M., Dorn, K.M., Chao, W.S., Watkin, E.E., Hernandez, A.G., Marks, M.D. and Gesch, R. (2018) Expression of FLOWERING LOCUS C and a frameshift mutation of this gene on chromosome 20 differentiate a summer and winter annual biotype of *Camelina sativa*. Plant Direct, 2, e00060.

Aubert, D., Chen, L., Moon, Y.-H., Martin, D., Castle, L.A., Yang, C.-H. and Sung, Z.R. (2001) EMF1, a novel protein involved in the control of shoot architecture and flowering in Arabidopsis. The Plant Cell, 13, 1865–1875.

Berti, M., Samarappuli, D., Johnson, B.L. and Gesch, R.W. (2017) Integrating winter camelina into maize and soybean cropping systems. Industrial crops and products, 107, 595–601.

Bolger, A.M., Lohse, M. and Usadel, B. (2014) Trimmomatic: A flexible trimmer for Illumina sequence data. Bioinformatics, 30, 2114–2120.

Bouché, F., Lobet, G., Tocquin, P. and Périlleux, C. (2016) FLOR-ID: an interactive database of flowering-time gene networks in Arabidopsis thaliana. Nucleic Acids Research, 44, D1167–D1171.

Brock, J.R., Donmez, A.A., Beilstein, M.A. and Olsen, K.M. (2018) Phylogenetics of *Camelina* Crantz. (Brassicaceae) and insights on the origin of gold-of-pleasure (Camelina sativa). Molecular Phylogenetics and Evolution 127, 834–842.

Broman, K.W., Wu, H., Sen, Ś. and Churchill, G.A. (2003) R/qtl: QTL mapping in experimental crosses. Bioinformatics, 19, 889–890.

Chao, W.S., Wang, H., Horvath, D.P. and Anderson, J.V. (2019) Selection of endogenous reference genes for qRT-PCR analysis in *Camelina sativa* and identification of *FLOWERING LOCUS C* allele-specific markers to differentiate summer-and winter-biotypes. Industrial crops and products, 129, 495–502.

Chaudhary, R., Koh, C.S., Kagale, S., Tang, L., Wu, S.W., Lv, Z., Mason, A.S., Sharpe, A.G., Diederichsen, A. and Parkin, I.A.P. (2020) Assessing Diversity in the *Camelina* Genus Provides Insights into the Genome Structure of *Camelina sativa*. G3: Genes, Genomes, Genetics, 10, 1297-1308.

Chaudhary, R., Koh, C.S., Perumal, S., Jin, L., Higgins, E.E., Kagale, S., Smith, M.A., Sharpe, A.G. and Parkin, I.A.P. (2022) Sequencing of *Camelina neglecta*, a diploid progenitor of the hexaploid oilseed *Camelina sativa*. Plant Biotechnology Journal.

Deng, W., Ying, H., Helliwell, C.A., Taylor, J.M., Peacock, W.J. and Dennis, E.S. (2011) *FLOWERING LOCUS C* (*FLC*) regulates development pathways throughout the life cycle of Arabidopsis. Proceedings of the National Academy of Sciences, 108, 6680–6685.

Dobin, A., Davis, C.A., Schlesinger, F., Drenkow, J., Zaleski, C., Jha, S., Batut, P., Chaisson, M. and Gingeras, T.R. (2013) STAR: Ultrafast universal RNA-seq aligner. Bioinformatics, 29, 15–21.

Fowler, S., Lee, K., Onouchi, H., Samach, A., Richardson, K., Morris, B., Coupland, G. and Putterill, J. (1999) *GIGANTEA*: a circadian clock-controlled gene that regulates photoperiodic flowering in Arabidopsis and encodes a protein with several possible membrane-spanning domains. The EMBO journal, 18, 4679–4688.

Galasso, I., Manca, A., Braglia, L., Ponzoni, E. and Breviario, D. (2015) Genomic fingerprinting of *Camelina* species using cTBP as molecular marker. American Journal of Plant Sciences, 6, 1184–1200.

Ge, S.X., Jung, D. and Yao, R. (2019) ShinyGO: a graphical gene-set enrichment tool for animals and plants. Bioinformatics, 36, 2628–2629.

Gehringer, A., Friedt, W., Lühs, W. and Snowdon, R.J. (2006) Genetic mapping of agronomic traits in false flax (*Camelina sativa* subsp. *sativa*). Genome, 49, 1555–1563.

Gel, B. and Serra, E. (2017) KaryoploteR: an R/Bioconductor package to plot customizable genomes displaying arbitrary data. Bioinformatics, 33, 3088–3090.

Gesch, R.W., Matthees, H.L., Alvarez, A.L. and Gardner, R.D. (2018) Winter camelina: Crop growth, seed yield, and quality response to cultivar and seeding rate. Crop Science, 58, 2089–2098.

Grabherr, M.G., Haas, B.J., Yassour, M., Levin, J.Z., Thompson, D.A., Amit, I., Adiconis, X., Fan, L., Raychowdhury, R. and Zeng, Q. (2011) Trinity: reconstructing a full-length transcriptome without a genome from RNA-Seq data. Nature biotechnology, 29, 644.

Hall, N. (2013) After the gold rush. Genome Biology, 14, 115.

Hansen, L.N. (1998) Intertribal somatic hybridization between rapid cycling *Brassica oleracea* L. and *Camelina sativa* (L.) Crantz. Euphytica, 104, 173–179.

Holtz, Y., David, J.L. and Ranwez, V. (2017) The genetic map comparator: a user-friendly application to display and compare genetic maps. Bioinformatics, 33.

Jiang, J.J., Zhao, X.X., Tian, W., Li, T.B. and Wang, Y.P. (2009) Intertribal somatic hybrids between *Brassica napus* and *Camelina sativa* with high linolenic acid content. Plant Cell, Tissue and Organ Culture, 99, 91–95.

Julié-Galau, S., Bellec, Y., Faure, J.D. and Tepfer, M. (2014) Evaluation of the potential for interspecific hybridization between *Camelina sativa* and related wild Brassicaceae in anticipation of field trials of GM camelina. Transgenic Research, 23, 67–74.

Kagale, S., Koh, C., Nixon, J., Bollina, V., Clarke, W.E., Tuteja, R., Spillane, C., Robinson, S.J., Links, M.G., Clarke, C., Higgins, E.E., Huebert, T., Sharpe, A.G. and Parkin, I.A.P. (2014) The emerging biofuel crop *Camelina sativa* retains a highly undifferentiated hexaploid genome structure. Nature Communications, 5, 1–11.

Kemi, U., Niittyvuopio, A., Toivainen, T., Pasanen, A., Quilot-Turion, B., Holm, K., Lagercrantz, U., Savolainen, O. and Kuittinen, H. (2013) Role of vernalization and of duplicated *FLOWERING LOCUS C* in the perennial *Arabidopsis lyrata*. New Phytologist, 197, 323–335.

Kim, D.-H., Doyle, M.R., Sung, S. and Amasino, R.M. (2009) Vernalization: winter and the timing of flowering in plants. Annual Review of Cell and Developmental, 25, 277–299.

King, K., Li, H., Kang, J. and Lu, C. (2019) Mapping quantitative trait loci for seed traits in *Camelina sativa*. Theoretical Applied Genetics, 132, 2567–2577.

Kurasiak-popowska, D., Graczyk, M. and Stuper-szablewska, K. (2020) Winter Camelina seeds as a raw material for the production of erucic acid-free oil. Food Chemistry, 127265.

Langfelder, P. and Horvath, S. (2008) WGCNA: an R package for weighted correlation network analysis. BMC Bioinformatics, 9, 559.

Li, B. and Dewey, C.N. (2011) RSEM: accurate transcript quantification from RNA-Seq data with or without a reference genome. BMC Bioinformatics, 12, 323.

Li, H. and Durbin, R. (2009) Fast and accurate short read alignment with Burrows-Wheeler transform. Bioinformatics (Oxford, England), 25, 1754–1760.

Li, H., Handsaker, B., Wysoker, A., Fennell, T., Ruan, J., Homer, N., Marth, G., Abecasis, G., Durbin, R. and Subgroup, G.P.D.P. (2009) The Sequence Alignment/Map format and SAMtools. Bioinformatics, 25, 2078–2079.

Li, H., Hu, X., Lovell, J.T., Grabowski, P.P., Mamidi, S., Chen, C., Amirebrahimi, M., Kahanda, I., Mumey, B. and Barry, K. (2021) Genetic dissection of natural variation in oilseed traits of camelina by whole-genome resequencing and QTL mapping. The plant genome, 14, e20110.

Lily, Z.L., Fahlgren, N., Kutchan, T., Schachtman, D., Ge, Y., Gesch, R., George, S., Dyer, J. and Abdel-Haleem, H. (2021) Discovering candidate genes related to flowering time in the spring panel of *Camelina sativa*. Industrial Crops and Products, 173, 114104.

Love, M.I., Huber, W. and Anders, S. (2014) Moderated estimation of fold change and dispersion for RNA-seq data with DESeq2. Genome Biology, 15, 550.

Luo, Z., Brock, J., Dyer, J.M., Kutchan, T.M., Augustin, M., Schachtman, D.P., Ge, Y., Fahlgren, N. and Abdel-Haleem, H. (2019a) Genetic diversity and population structure of a *Camelina sativa* spring panel. Frontiers in Plant Science, 10, 184.

Luo, Z., Tomasi, P., Fahlgren, N. and Abdel-Haleem, H. (2019b) Genome-wide association study (GWAS) of leaf cuticular wax components in *Camelina sativa* identifies genetic loci related to intracellular wax transport. BMC Plant Biology, 19, 187.

Madeira, F., Pearce, M., Tivey, A.R., Basutkar, P., Lee, J., Edbali, O., Madhusoodanan, N., Kolesnikov, A. and Lopez, R. (2022) Search and sequence analysis tools services from EMBL-EBI in 2022. Nucleic acids research, 50, W276–W279.

Martin, S.L., Lujan-Toro, B.E., Sauder, C.A., James, T., Ohadi, S. and Hall, L.M. (2019) Hybridization rate and hybrid fitness for *Camelina microcarpa* Andrz. ex DC (♀) and *Camelina sativa* (L.) Crantz(Brassicaceae) (♂). Evolutionary Applications, 12, 443–455.

Martin, S.L., Sauder, C.A., James, T., Cheung, K.W., Razeq, F.M., Kron, P. and Hall, L. (2015) Sexual hybridization between *Capsella bursa-pastoris* (L.) Medik (♀) and *Camelina sativa* (L.) Crantz (♂) (Brassicaceae). Plant Breeding, 134, 212–220.

Martin, S.L., Smith, T.W., James, T., Shalabi, F., Kron, P. and Sauder, C.A. (2017) An update to the Canadian range, abundance, and ploidy of *Camelina* spp. (Brassicaceae) east of the Rocky Mountains. Botany, 95, 405–417.

McKenna, A., Hanna, M., Banks, E., Sivachenko, A., Cibulskis, K., Kernytsky, A., Garimella, K., Altshuler, D., Gabriel, S., Daly, M. and DePristo, M.A. (2010) The Genome Analysis Toolkit: A MapReduce framework for analyzing next-generation DNA sequencing data. Genome Research, 20, 1297–1303.

Michaels, S.D. and Amasino, R.M. (1999) *FLOWERING LOCUS C* encodes a novel MADS domain protein that acts as a repressor of flowering. The Plant Cell, 11, 949–956.

Mizoguchi, T., Wright, L., Fujiwara, S., Cremer, F.d.r., Lee, K., Onouchi, H., Mouradov, A., Fowler, S., Kamada, H. and Putterill, J. (2005) Distinct roles of GIGANTEA in promoting flowering and regulating circadian rhythms in Arabidopsis. The Plant Cell, 17, 2255–2270.

Mouradov, A., Cremer, F. and Coupland, G. (2002) Control of flowering time: interacting pathways as a basis for diversity. The Plant Cell, 14, S111–S130.

Narasimhulu, S.B., Kirti, P.B., Bhatt, S.R., Prakash, S. and Chopra, V.L. (1994) Intergeneric protoplast fusion between *Brassica carinata* and *Camelina sativa*. Plant Cell Reports, 13, 657–660.

O’Neill, C.M., Lu, X., Calderwood, A., Tudor, E.H., Robinson, P., Wells, R., Morris, R. and Penfield, S. (2019) Vernalization and floral transition in autumn drive winter annual life history in oilseed rape. Current Biology, 29, 4300–4306. e4302.

Okazaki, K., Sakamoto, K., Kikuchi, R., Saito, A., Togashi, E., Kuginuki, Y., Matsumoto, S. and Hirai, M.J.T. (2007) Mapping and characterization of FLC homologs and QTL analysis of flowering time in *Brassica oleracea*. Theoretical Applied Genetics, 114, 595–608.

Poland, J.A., Brown, P.J., Sorrells, M.E. and Jannink, J.-L. (2012) Development of high-density genetic maps for barley and wheat using a novel two-enzyme genotyping-by-sequencing approach. PloS One, 7, e32253.

Sasaki, E., Zhang, P., Atwell, S., Meng, D. and Nordborg, M. (2015) " Missing" G x E variation controls flowering time in *Arabidopsis thaliana*. PLoS Genet, 11, e1005597.

Schiessl, S.V., Quezada-Martinez, D., Tebartz, E., Snowdon, R.J. and Qian, L. (2019) The vernalisation regulator *FLOWERING LOCUS C* is differentially expressed in biennial and annual *Brassica napus*. Scientific Reports, 9, 1–15.

Séguin-Swartz, G., Nettleton, J.A., Sauder, C., Warwick, S.I. and Gugel, R.K. (2013) Hybridization between *Camelina sativa* (L.) Crantz (false flax) and North American *Camelina* species. Plant Breeding, 132, 390–396.

Sheldon, C.C., Rouse, D.T., Finnegan, E.J., Peacock, W.J. and Dennis, E.S. (2000) The molecular basis of vernalization: the central role of *FLOWERING LOCUS C* (*FLC*). Proceedings of the National Academy of Sciences, 97, 3753–3758.

Simpson, G.G., Dijkwel, P.P., Quesada, V., Henderson, I. and Dean, C. (2003) FY is an RNA 3′ end-processing factor that interacts with FCA to control the Arabidopsis floral transition. Cell, 113, 777–787.

Singh, R., Bollina, V., Higgins, E.E., Clarke, W.E., Eynck, C., Sidebottom, C., Gugel, R., Snowdon, R. and Parkin, I.A.P. (2015) Single-nucleotide polymorphism identification and genotyping in *Camelina sativa*. Molecular Breeding, 35, 35.

Stringam, G.R. (1971) Genetics of four hypocotyl mutants in Brassica campestris L. Journal of Heredity, 62, 248–250.

Swiezewski, S., Liu, F., Magusin, A. and Dean, C. (2009) Cold-induced silencing by long antisense transcripts of an Arabidopsis Polycomb target. Nature Genetics, 462, 799.

Takada, S., Akter, A., Itabashi, E., Nishida, N., Shea, D.J., Miyaji, N., Mehraj, H., Osabe, K., Shimizu, M. and Takasaki-Yasuda, T. (2019) The role of FRIGIDA and FLOWERING LOCUS C genes in flowering time of *Brassica rapa* leafy vegetables. Scientific Reports, 9, 1–11.

Team, R.C. (2021) R: A language and environment for statistical computing. R Foundation for Statistical Computing, Vienna, Austria. URL https://www.R-project.org.

Teotia, S. and Tang, G. (2015) To bloom or not to bloom: role of microRNAs in plant flowering. Molecular plant, 8, 359–377.

Thorvaldsdóttir, H., Robinson, J.T. and Mesirov, J.P. (2012) Integrative Genomics Viewer (IGV): high-performance genomics data visualization and exploration. Briefings in Bioinformatics, 14, 178–192.

Vollmann, J. and Eynck, C. (2015) Camelina as a sustainable oilseed crop: Contributions of plant breeding and genetic engineering. Biotechnology Journal, 10, 525–535.

Vollmann, J., Grausgruber, H., Stift, G., Dryzhyruk, V. and Lelley, T. (2005) Genetic diversity in camelina germplasm as revealed by seed quality characteristics and RAPD polymorphism. Plant Breeding, 124, 446–453.

Voorrips, R. (2002) MapChart: software for the graphical presentation of linkage maps and QTLs. Journal of Heredity, 93, 77–78.

Wickham, H. (2016) ggplot2: elegant graphics for data analysis: springer.

Wittenberg, A., Anderson, J.V. and Berti, M.T. (2019) Winter and summer annual biotypes of camelina have different morphology and seed characteristics. Industrial Crops and Products, 135, 230–237.

Wu, Y., Bhat, P., Close, T.J. and Lonardi, S. (2007) Efficient and accurate construction of genetic linkage maps from noisy and missing genotyping data. In International Workshop on Algorithms in Bioinformatics: Springer, pp. 395-406.

Xiao, D., Zhao, J.J., Hou, X.L., Basnet, R.K., Carpio, D.P., Zhang, N.W., Bucher, J., Lin, K., Cheng, F. and Wang, X.W. (2013) The *Brassica rapa* FLC homologue FLC2 is a key regulator of flowering time, identified through transcriptional co-expression networks. Journal of Experimental Botany, 64, 4503–4516.

Zhao, J., Kulkarni, V., Liu, N., Pino Del Carpio, D., Bucher, J. and Bonnema, G. (2010) BrFLC2 (FLOWERING LOCUS C) as a candidate gene for a vernalization response QTL in *Brassica rapa*. Journal of Experimental Botany, 61, 1817–1825.

Zhou, Y., Zhou, B., Pache, L., Chang, M., Khodabakhshi, A.H., Tanaseichuk, O., Benner, C. and Chanda, S.K. (2019) Metascape provides a biologist-oriented resource for the analysis of systems-level datasets. Nature communications, 10, 1-10.

Zhu, A., Ibrahim, J.G. and Love, M.I. (2018) Heavy-tailed prior distributions for sequence count data: removing the noise and preserving large differences. Bioinformatics, 35, 2084–2092.

Zou, X., Suppanz, I., Raman, H., Hou, J., Wang, J., Long, Y., Jung, C. and Meng, J. (2012) Comparative analysis of FLC homologues in Brassicaceae provides insight into their role in the evolution of oilseed rape. PloS One, 7, e45751.

