## Supplementary Figures for "Mapping QTL for vernalization requirement identified adaptive divergence of the candidate gene *Flowering Locus C* in polyploid *Camelina sativa*"

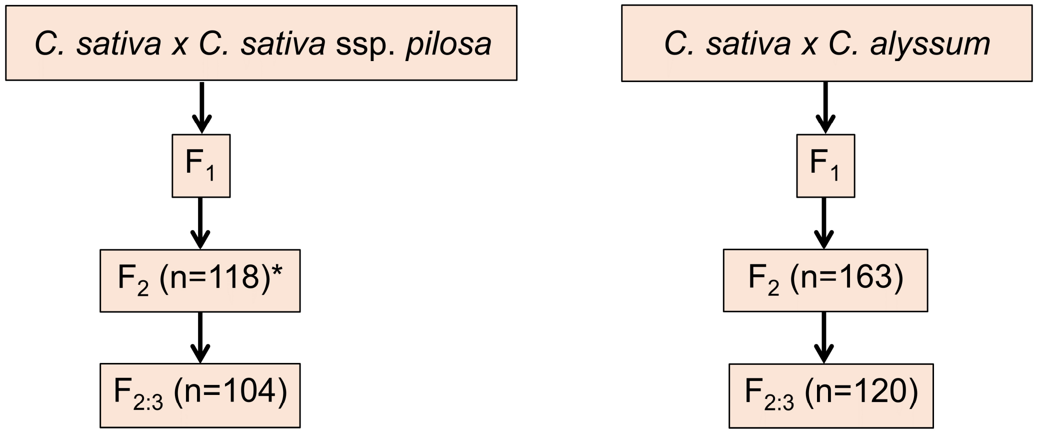


**Figure S1. Inter- and intraspecific hybridization scheme adopted in this study with total number of lines for segregating populations.** Accessions TMP23992 (*C. sativa*), CN113692 (*C. sativa* ssp. *pilosa*) and CAM176 (*C. alyssum*) were used in this study. $F_1$ was self-pollinated to produce $F_2$ lines and a single seed from individual $F_2$ lines were used to generate $F_{2:3}$ lines. The total number of lines phenotyped at each generation are shown in parenthesis.

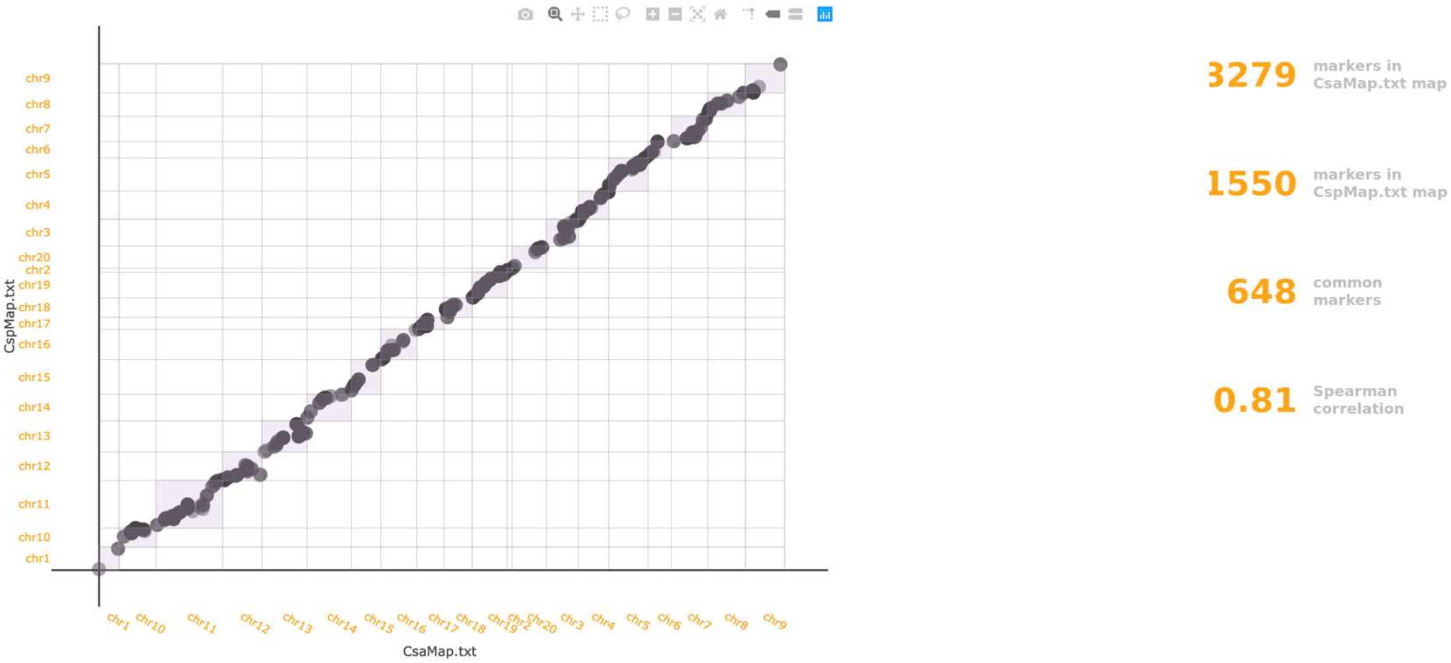


**Figure S2. A collinearity between the Csa (*C. sativa* × *C. alyssum*) Csp (*C. sativa* × *C. sativa* ssp. *pilosa*) genetic maps using The genetic Map Comparator** (https://bioweb.supagro.inrae.fr/geneticMapComparator/).

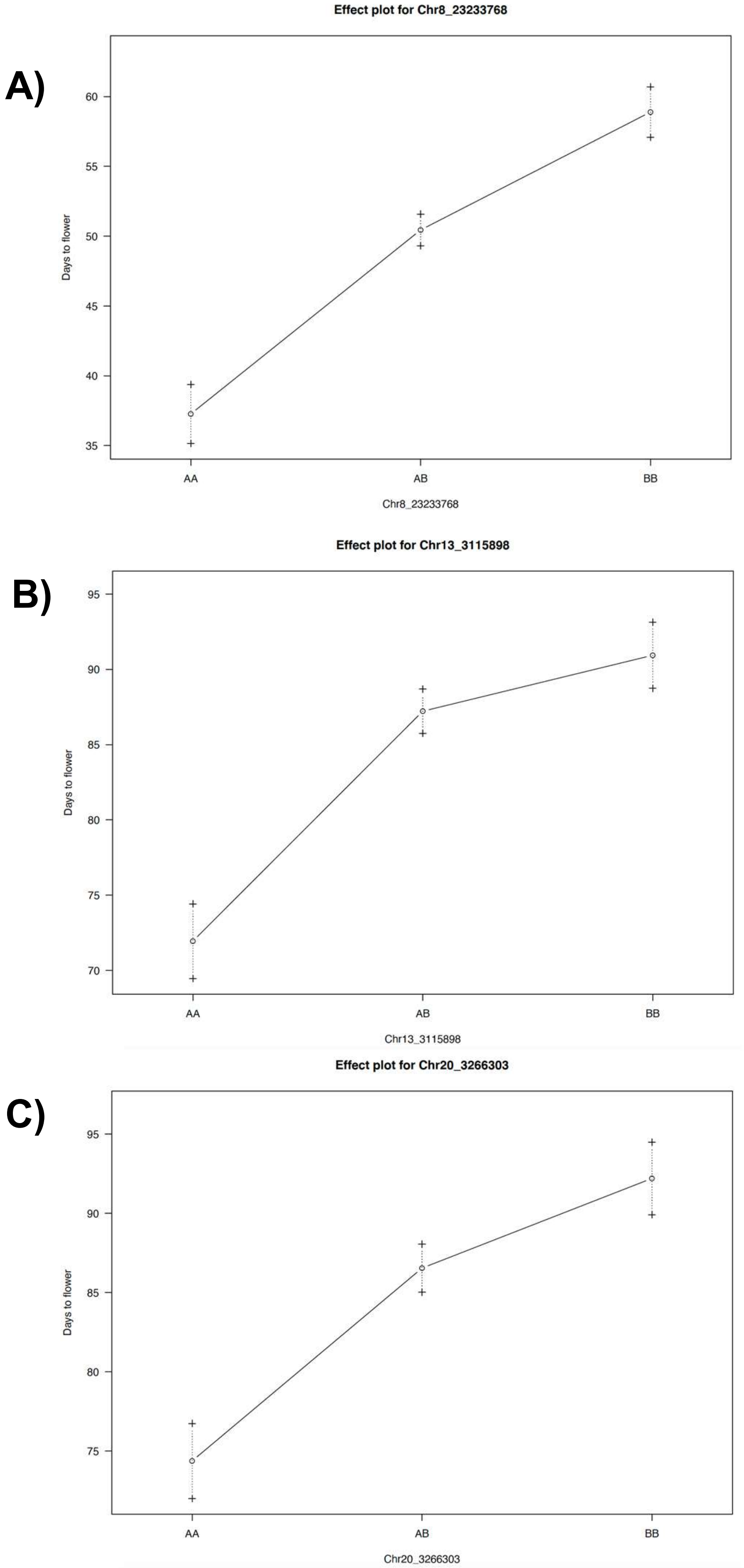


**Figure S3. Effect of parental alleles on days to flower.** A) QTL region on chr 8 from Csp (*C. sativa* × *C. sativa* ssp. *pilosa*) population. B) QTL region on chromosome 13 from Csa (*C. sativa* × *C. alyssum*) population, and C) QTL region on chromosome 20 from Csa population. A - represents spring-type allele, B - winter-type allele.

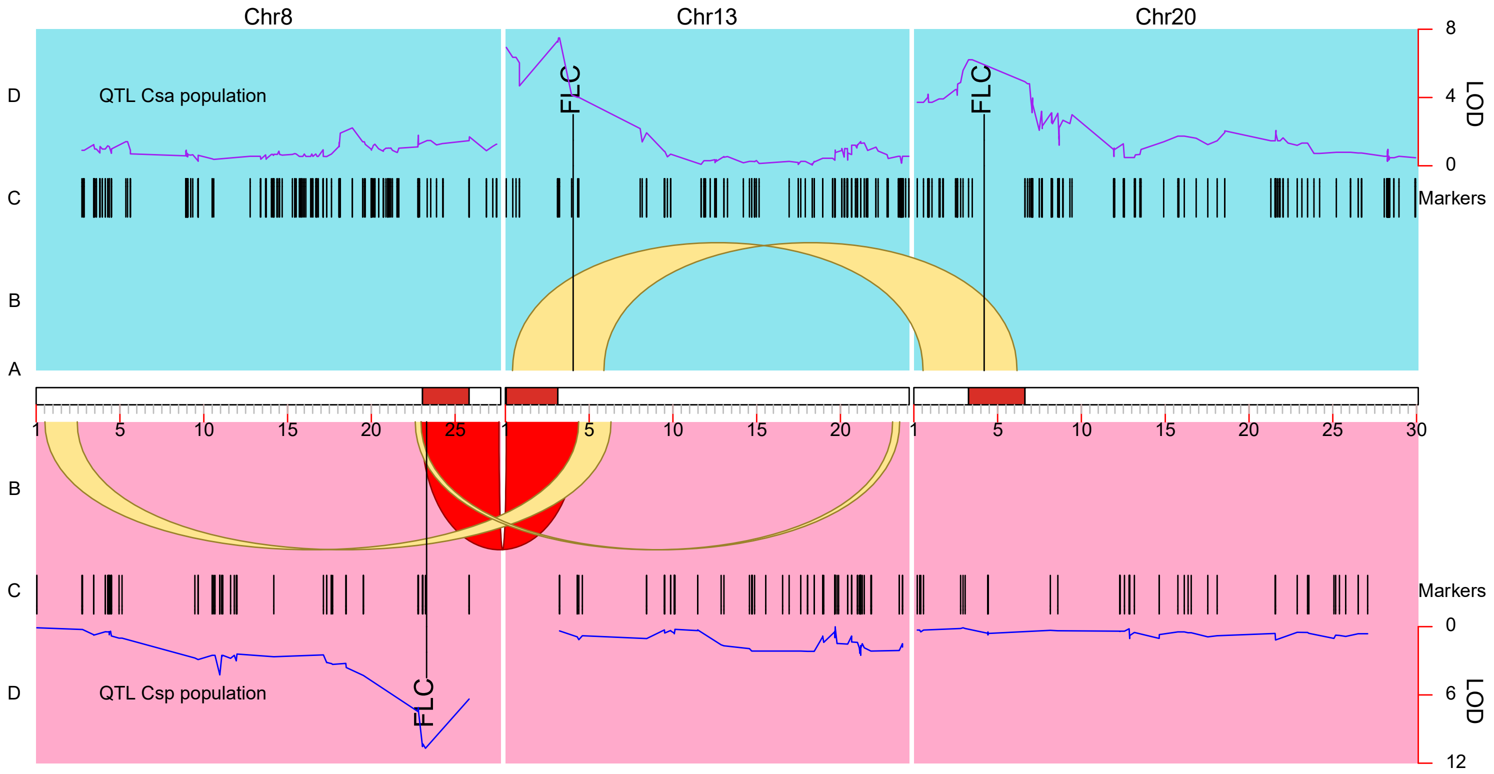


**Figure S4. Structure of QTL regions for vernalization requirement in *C. sativa*.** A) Three homoeologous chromosomes are represented with the QTL confidence interval in red and position of *FLC* gene shown above for Csa (*C. sativa* × *C. alyssum*) and below for Csp (*C. sativa* × *C. sativa* ssp. *pilosa*) ; B) Yellow ribbons represent syntenic genes around the QTL among homoeologous chromosomes; red ribbon represents an inversion, C) Distribution of markers from Csa (top) and Csp (bottom), and D) Distribution of LOD score for identified QTL in Csa (top) and Csp (bottom) populations.

(A`

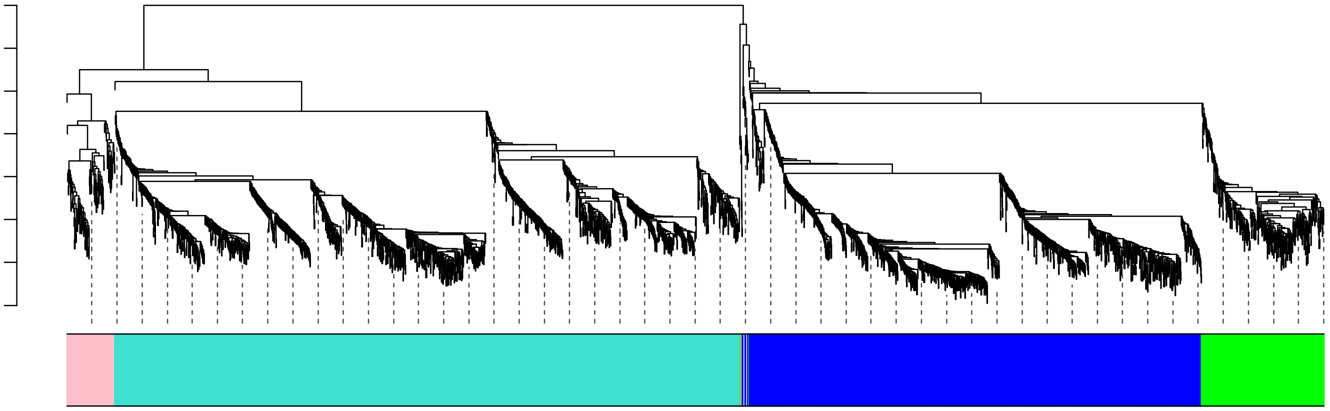

(B`

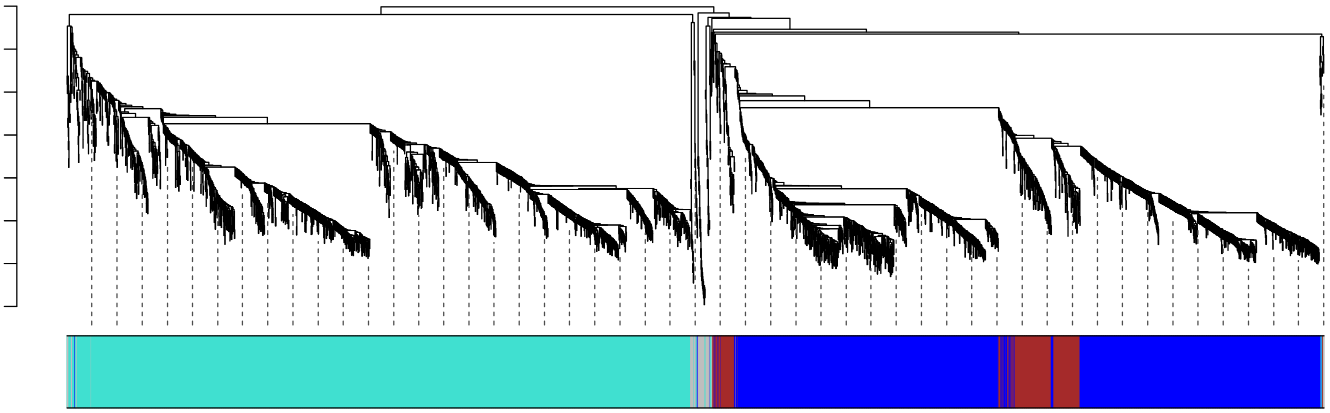

(C`

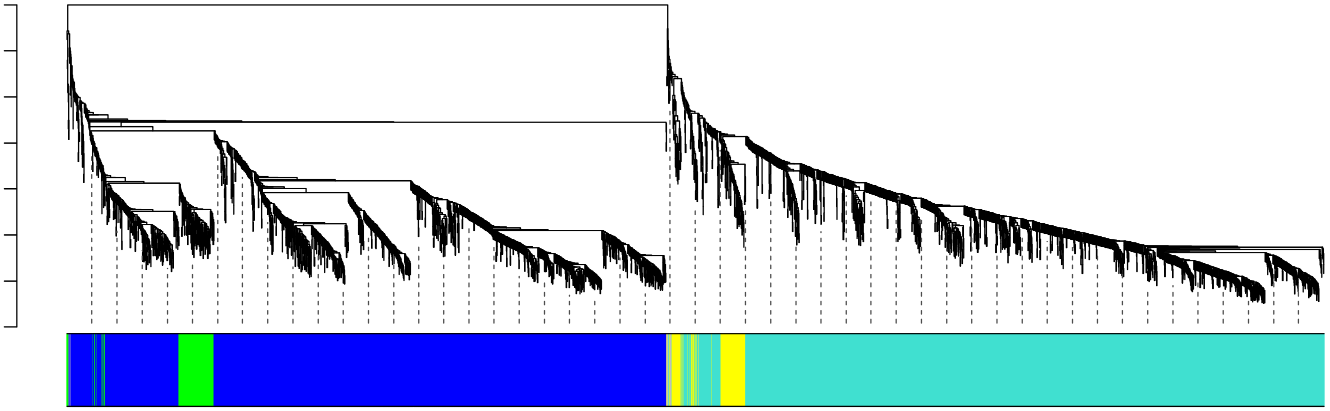

**Figure S5. Gene co-expression analysis using WGCNA.** Branches in the hierarchical clustering dendrograms corresponds to module where a different colour represents a different module for TMP23992 (A), CN113692 (B), and CAM176 (C).

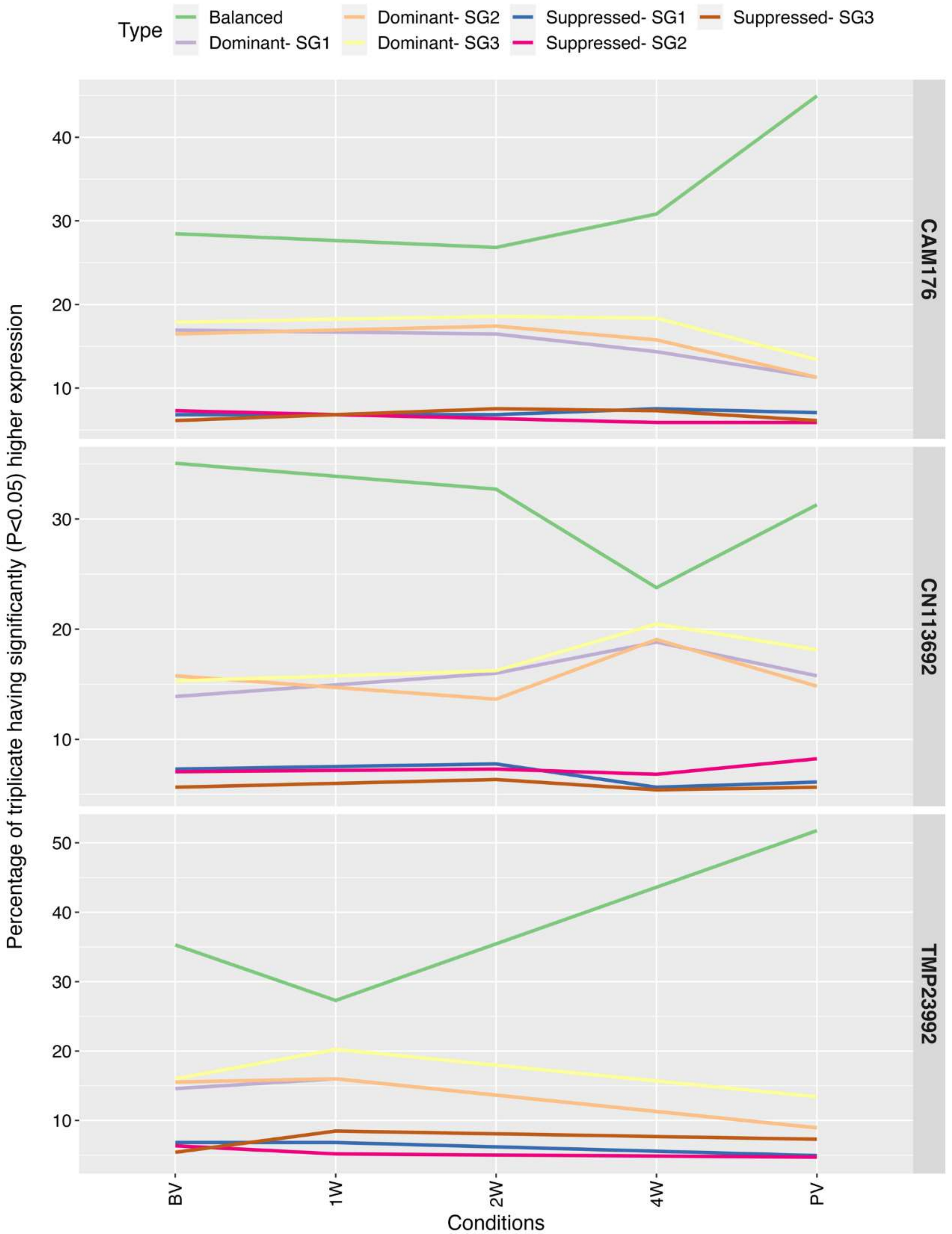


**Figure S6. Subgenome dominance analysis with flowering related genes in Camelina species.** The expression dominance for a triplicate gene were compared across different genotypes CAM176 , CN113692, and TMP23992; and different time points. BV is before-vernalization, 1W, 2W, and 4W are one, two or four weeks during vernalization, and PV is one week post-vernalization.

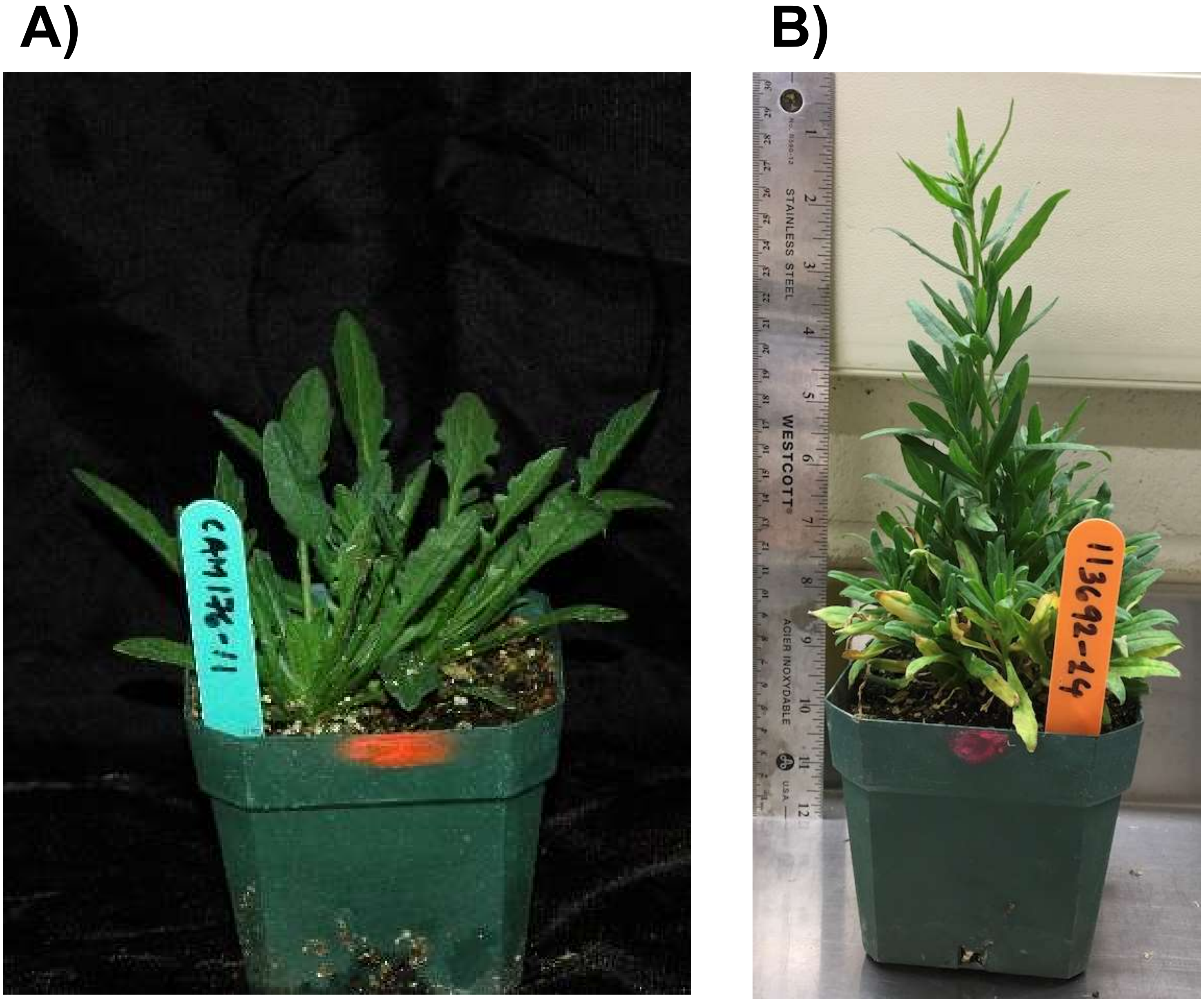


**Figure S7. Plant growth in two winter-type Camelina lines without vernalization.** A) *C. alyssum* shows reduced stem and profuse leaves (30 days after sowing); and B) *C. sativa* ssp. *pilosa* (60 days after sowing without vernalization) characterised by stem elongation and branching.

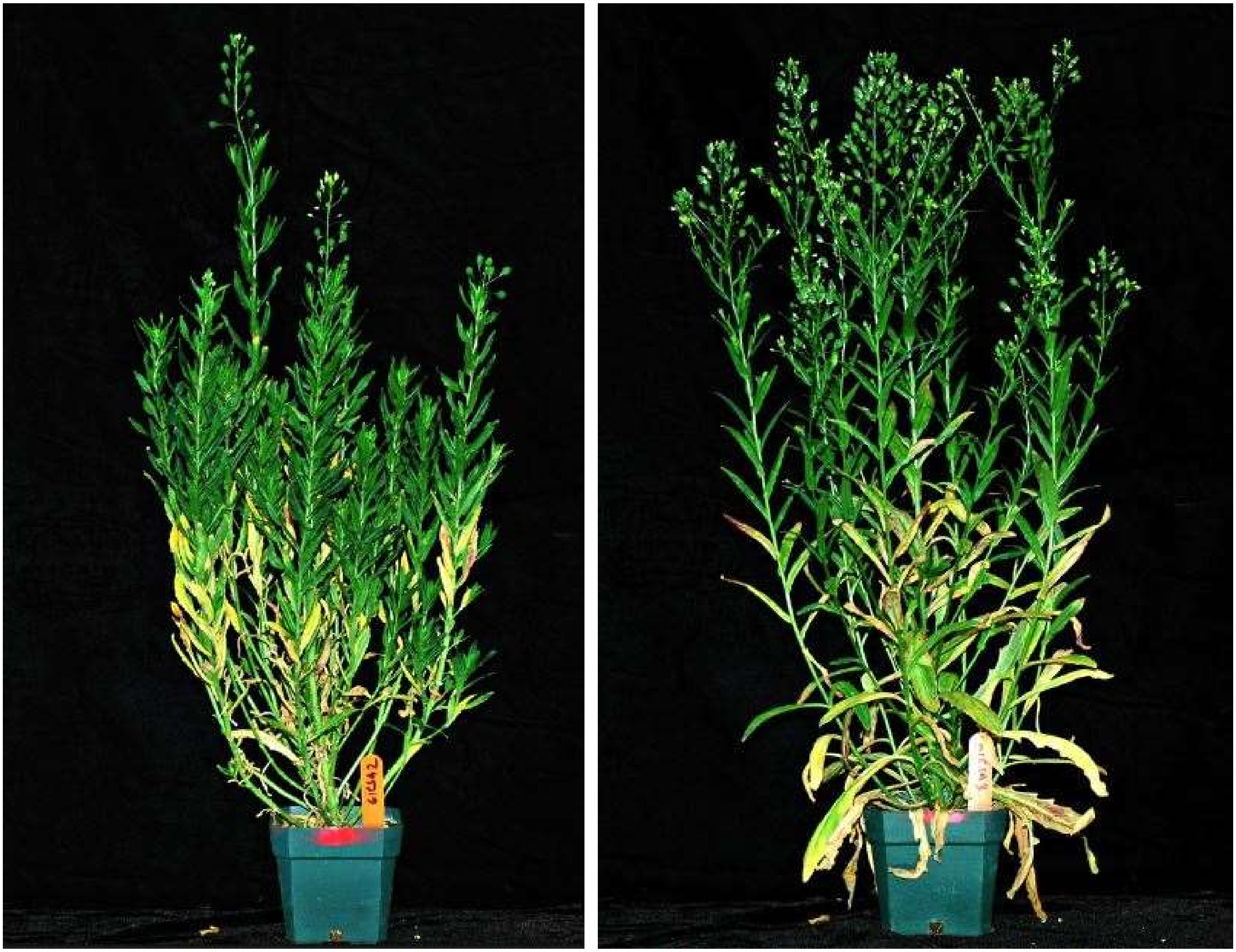

**Figure S8. Effect of duration of vernalization on flowering in hybrid developed from *C. sativa* × *C. alyssum* after 1 week (left) and three weeks (right) into vernalization.** Flowering at particular branch and rest branches are in vegetative stage (95 days after seeding).

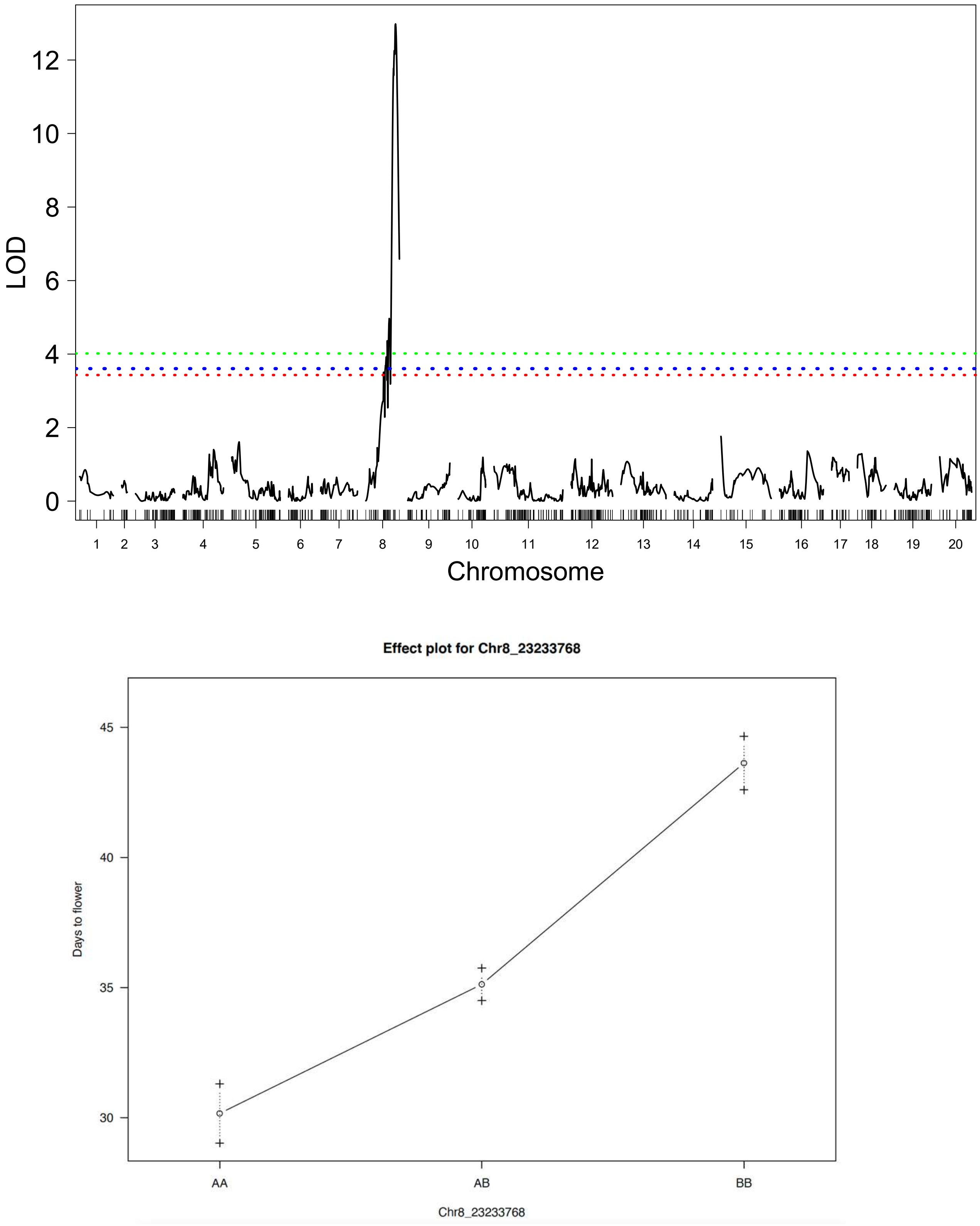


**Figure S9. QTL mapping for days to flower from $F_{2:3}$ derived from Csp (*C. sativa* × *C. sativa* ssp. *pilosa*) population.** A) LOD Score distribution across the chromosomes; B) Effect of parental allele at QTL locus, A - spring-type allele, B - winter-type allele.

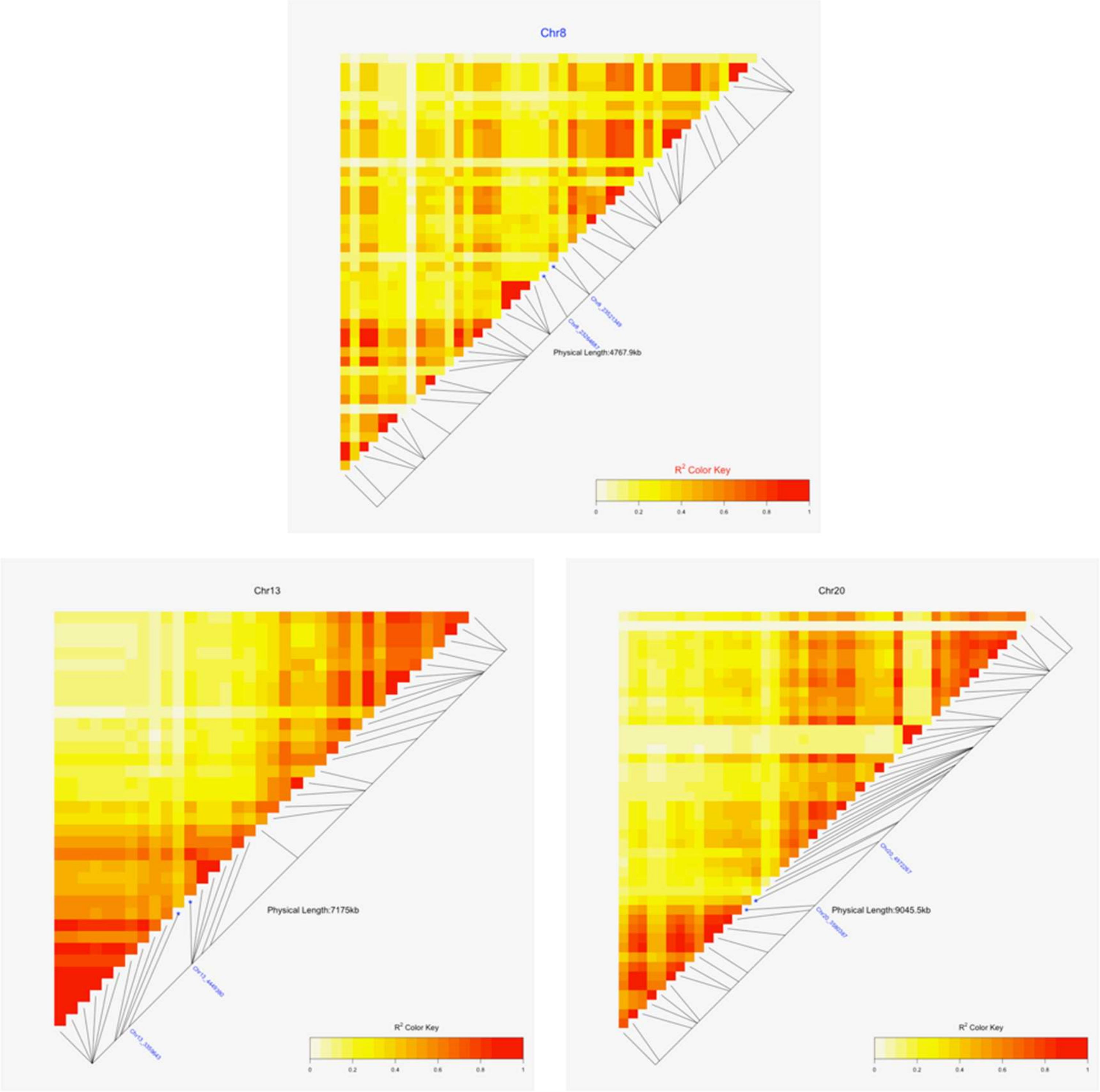


**Figure S10 Linkage disequilibrium heatmap showing $R^2$ for markers across the QTL regions on chromosomes 8, 13 and 20.** The markers labelled in blue are those flanking *FLC.*

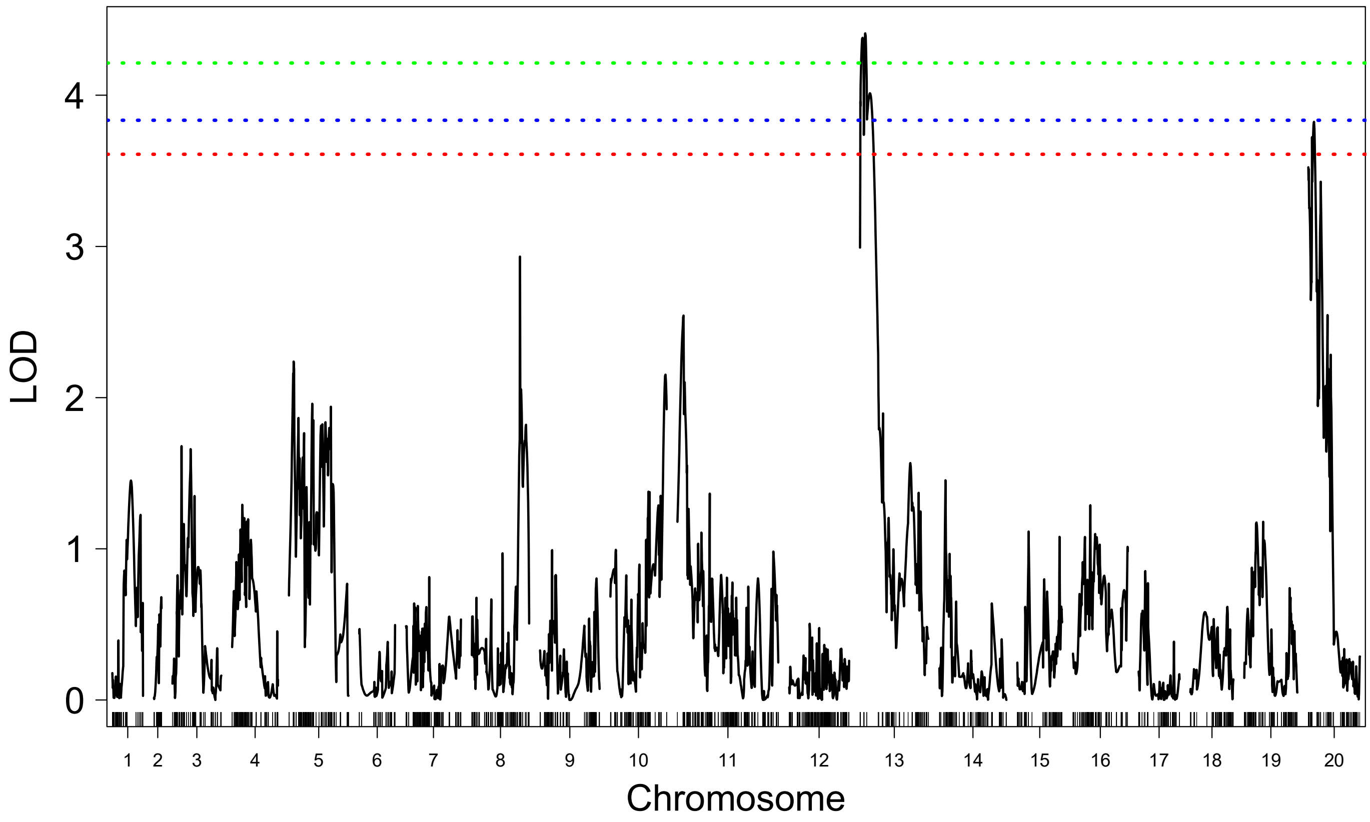


```
                 fitqtl summary

Method: Haley-Knott regression
Model:  normal phenotype
Number of observations : 161

Full model result
----------------------------------
Model formula: y ~ Q1 + Q2

       df       SS        MS      LOD     %var Pvalue(Chi2)    Pvalue(F)
Model   4 19214.46 4803.6144 8.286923 21.10366 1.037217e-07 1.631174e-07
Error 156 71833.53  460.4713
Total 160 91047.99


Drop one QTL at a time ANOVA table:
----------------------------------
        df Type III SS   LOD   %var F value Pvalue(Chi2) Pvalue(F)
13@12.0  2        9830 4.484 10.796  10.674            0  4.52e-05 ***
20@12.0  2        8462 3.893  9.294   9.188            0  0.000169 ***
---
Signif. codes:  0 ‘***’ 0.001 ‘**’ 0.01 ‘*’ 0.05 ‘.’ 0.1 ‘ ’ 1
```

**Figure S11. QTL mapping of days to flowering from $F_{2:3}$ derived from Csa (*C. sativa* × *C. alyssum*) population.**

```
CLUSTAL O(1.2.4) multiple sequence alignment


CAM176_chr20          AQFQAKKEEKRLVSLRRLEPNPNLRIKLGHKGLSETEAMGRKKLEIKRIENKSSRQVTFS      60
DH55_chr20            ------KEEKRLVSLRRLEPNPNLRIKLGHKGLSETEAMGRKKLEIKRIENKSSRQVTFS      54
TMP23992_chr20        --------------------------------------MGRKKLEIKRIENKSSRQVTFS      22
DH55_chr8             --------------------------------------MGRKKLEIKRIENKSSRQVTFS      22
TMP23992_chr8         --------------------------------------MGRKKLEIKRIENKSSRQVTFS      22
DH55_chr13            --------------------------------------MGRKKLEIKRIENKSSRQVTFS      22
CAM176_chr13          --------------------------------------MGRKKLEIKRIENKSSRQVTFS      22
Cneglecta_chr05       --------------------------------------MGRKKLEIKRIENKSSRQVTFS      22
CN113692_chr8         --------------------------------------MGRKKLEIKRIENKSSRQVTFS      22
CAM176_chr8           --------------------------------------MGRKKLEIKRIENKSSRQVTFS      22
                                                            **********************

CAM176_chr20          KRRNGLIEKARQLSVLCDASVALLVVSASGKLYSFSSGDNLVKILDRYGKQHADDLKALD      120
DH55_chr20            KRRNGLIEKARQLSVLCDASVALLVVSASGKLYSFSSGDNLVKILDRYGKQHADDLKALD      114
TMP23992_chr20        KRRNGLIEKARQLSVLCDASVALLVVSASGKLYSFSSGDNLVKILDRYGKQHADDLKALD      82
DH55_chr8             KRRNGLIEKARQLSVLCDASVALLVVSASGKLYSFSSGDNLVKILDRYGKQHADDLKALD      82
TMP23992_chr8         KRRNGLIEKARQLSVLCDASVALLVVSASGKLYSFSSGDNLVKILDRYGKQHADDLKALD      82
DH55_chr13            KRRNGLIEKARQLSVLCDASVALLVVSASGKLYSFSSGDNLVKILDRYGKQHADDLKALD      82
CAM176_chr13          KRRNGLIEKARQLSVLCDASVALLVVSASGKLYSFSSGDNLVKILDRYGKQHADDLKALD      82
Cneglecta_chr05       KRRNGLIEKARQLSVLCDASVALLVVSASGKLYSFSSGDNLVKILDRYGKQHADDLKALD      82
CN113692_chr8         KRRNGLIEKARQLSVLCDASVALLVVSASGKLYSFSSGDNLVKILDRYGKQHADDLKALD      82
CAM176_chr8           KRRNGLIEKARQLSVLCDASVALLVVSASGKLYSFSSGDNLVKILDRYGKQHADDLKALD      82
                      ************************************************************

CAM176_chr20          LQSKELNYGSHHELLELVESNLVESNVNNVSVDSLVQLEEHLETALSITRAKKTELMLKL      180
DH55_chr20            LQSKELNYGSHHELLELVESNLVESNVNNVSVDSLVQLEEHLETALSITRAKKTELM*--      171
TMP23992_chr20        LQSKELNYGSHHELLELVESNLVESNVNNVSVDSLVQLEEHLETALSITRAKKTELM*--      139
DH55_chr8             LQSKALNYGSHHELLELVESNLVESNVNNVSVDALV-LEEHLETALSVTSAKKTELMLKL      141
TMP23992_chr8         LQSKALNYGSHHELLELVESNLVESNVNNVSVDALV-LEEHLETALSVTSAKKTELMLKL      141
DH55_chr13            LQSKALNYGSHHELLELVESNLVESNVNNVSVDDLVQLEEHLETALSVTRAKKTELILKL      142
CAM176_chr13          LQSKALNYGSHHELLELVESNLVESNVNNVSVDDLVQLEEHLETALSVTRAKKTELILKL      142
Cneglecta_chr05       LQSKALNYGSHHELLELVESNLVESNVNNVSVDALVQLEEHLETALSVTRAKKTELMLKL      142
CN113692_chr8         LQSKALNYGSHHELLELVESNLVESNVNNVSVDALVQLEEHLETALSVTRAKKTELMLKL      142
CAM176_chr8           LQSKALNYGSHHELLELVESNLVESNVNNVSVDALVQLEEHLETALSVTRAKKTELMLKL      142
                      **** **************************** ** **********:* ******:

CAM176_chr20          VENLKEKEKLLKEENQVLASQMETNHVVGAEADMEMEMSPAGQISDNLPVTLPLLN*  236
DH55_chr20            ---------------------------------------------------------  171
TMP23992_chr20        ---------------------------------------------------------  139
DH55_chr8             VENLKEKEKLLKEENQVLASQMETNHVVGAEADMEMEMSPVGQISDNLPVTLPLLN*  197
TMP23992_chr8         VENLKEKEKLLKEENQVLASQMETNHVVGAEADMEMEMSPVGQISDNLPVTLPLLN*  197
DH55_chr13            VENLKEKEKLLKEENQVLARQMETNHVVGAEADMEMEMSPAGQISDNLPVTLPLLN*  198
CAM176_chr13          VENLKEKEKLLKEENQVLARQMETNHVVGAEADMEMEMSPAGQISDNLPVTLPLLN*  198
Cneglecta_chr05       VENLKEKEKLLKEENQVLASQMETNHVVGAEADMEMEMSPAGQISDNLPVTLPLLN*  198
CN113692_chr8         VENLKEKEKLLKEENQVLASQMETNHVVGAEADMEMEMSPVGQISDNLPVTLPLLN*  198
CAM176_chr8           VENLKEKEKLLKEENQVLASQMETNHVVGAEADMEMEMSPVGRISDNLPVTLPLLN*  198
```

**Figure S12. Reconstruction of *Flowering Locus C* (*FLC*) genes using Trinity from TMP23992 (*C. sativa*), DH55 (*C. sativa*), CN113692 (*C. sativa* ssp. *pilosa*), and CAM176 (*C. alyssum*) and comparison with *FLC* orthologs from DH55 (reference genome) and *C. neglecta* (reference genome).** Due to low gene expression, *FLC* on chromosome 13 and chromosome 20 for CN113692 and chromosome 13 for TMP23992 were not constructed.
